# IL-25-induced memory ILC2s mediate long-term small intestinal adaptation

**DOI:** 10.1101/2025.03.25.645270

**Authors:** Victor S. Cortez, Sara Viragova, Satoshi Koga, Meizi Liu, Claire E. O’Leary, Roberto R. Ricardo-Gonzalez, Andrew W. Schroeder, Nathan Kochhar, Ophir D. Klein, Michael S. Diamond, Hong-Erh Liang, Richard M. Locksley

**Author notes:** Current address: Immunology Frontier Research Center, Osaka, Japan (Satoshi Koga). **Current address: Department of Pediatrics, University of Wisconsin School of Medicine and Public Health, Madison, WI, USA. Corresponding Author/Lead Contact: Dr. Richard M. Locksley, University of California, San Francisco, 513 Parnassus Avenue, S-1032B, San Francisco, CA 94143-0795.

## Abstract

The adaptation of intestinal helminths to vertebrates evolved strategies to attenuate host tissue damage to support reproductive needs of parasites necessary to disseminate offspring to the environment. Helminths initiate the IL-25-mediated tuft cell-ILC2 circuit that enhances barrier protection of the host although viable parasites can target and limit the pathway. We used IL-25 to create small intestinal adaptation marked by anatomic, cell compositional and immunologic changes that persisted months after induction. Small intestinal adaptation was associated with heightened resistance to barrier pathogens, including in the lung, and sustained by transcriptionally and epigenetically modified, tissue-resident, memory-effector ILC2s distinct from those described by innate ‘training’; epithelial stem cells remained unaltered. Despite requiring IL-25 for induction, memory ILC2s maintained an activated state in the absence of multiple alarmins and supported mucosal resilience while avoiding adverse sensitization to chronic inflammation, revealing a pathway for deploying innate immune cells to coordinate a distributed mucosal defense.

## Introduction

Tissue adaptation through life reflects the accumulation of repeated stresses in cells that drive transcriptional, epigenetic and compositional changes that sustain homeostasis by promoting more rapid responses to injury and infection (1). While prior studies have emphasized roles for immune cells in adaptation to promote physiologically resilient states that sustain tissue homeostasis (2,3), it is unclear whether tissue adaptation is mediated collectively by the cells of a given organ or is hierarchically organized by key cell types. Although changes in resident immune cell populations commonly occur in tissues that adapt to inflammatory perturbation, recent studies have called attention to stable transcriptional and epigenetic changes in epithelial stem cells themselves, particularly in barrier tissues, that enable developmental and functional plasticity among epithelial progeny to promote rapid recovery from injury but at potential cost for hyperproliferative and inflammatory disorders (4,5).

We examined adaptation by the small intestine, a key barrier where efficient nutrient extraction requires a balance between immune tolerance to food and microbial commensals but resistance to toxins and pathogens as driven by continuous exposure to the external environment. Among undomesticated vertebrates and invertebrates, parasitic helminths are near ubiquitous residents of this intestinal interface, revealing a highly evolved relationship between host and helminth (6). Unlike bacteria, fungi and viruses, parasitic worms do not typically divide in their hosts but rather produce large numbers of offspring that are expelled in feces and mature in the environment. Immunity is acquired slowly and is typically incomplete, as characterized by combinations of tolerance, immune regulation, and concomitant immunity, the process whereby infested individuals gain resistance to additional infection by eggs or larvae despite the inability to clear fecund adults (7,8), including at distal sites like the lung (9). We examined the small intestinal response as sculpted by the evolutionary constraints on tissue damage by helminths to try to identify pathways that engage host protective rather than pathologic adaptation (10).

Over the past several years, we and others described the small intestine tuft cell – group 2 innate lymphoid cell (ILC2) circuit that is activated in response to certain luminal protists and helminths (11–13). In brief, rare chemosensory tuft cells express G-protein-coupled receptors (GPCRs) that recognize metabolites of luminal organisms and, when activated, release IL-25 and stimulate resident ILC2s which constitutively express the IL-25 receptor. ILC2s in turn release IL-13, which acts to bias differentiation of intestinal epithelia to secretory cells, including goblet and tuft cells, thus enhancing the protective mucus layer and the ‘sensing’ capacity at the epithelial interface. Chronic parasitism or genetic deletion of intrinsic inhibitory feedback pathways in ILC2s drives circuit activation leading to anatomic changes including small intestinal lengthening, changes in the cellular composition of epithelial and immune cells, metabolic stability, and infection resistance, but the stability of this adaptation and the cells and pathways necessary to sustain it remain undefined (14). Here, we used recombinant IL-25 to establish a prolonged state of small intestinal adaptation driven by a previously uncharacterized population of long-lived ILC2s; the epithelial stem cell compartment was unaltered. In contrast to the consensus definition of innate ‘trained’ immunity (15), by which ILCs become epigenetically modified and return to the basal state but increase their response upon secondary challenge, IL-25-induced ILC2s resembled memory-effector lymphocytes that accumulated and differentiated to a long-lasting activated phenotype sustained even in the absence of alarmin cytokines required to initiate the memory state.

## Results

### Induction of small intestinal adaptation by IL-25

Based on prior studies (14), we optimized a regimen of IL-25 injections to ask whether this pathway alone could induce persistent small intestinal anatomic and compositional adaptations resembling those caused by viable helminths. Mice were given either intraperitoneal recombinant IL-25 (500 ng) or the same volume of PBS (control) 3 times per week for four weeks and rested for at least 50 days prior to analysis (typically +50 to +75 unless otherwise specified). We designated +50 days the memory time point for most analyses and representing >10 complete turnovers of the intestinal epithelium that is only further increased by activation of type 2 immunity (16,17) (**Figure 1A**). IL-25-treated mice exhibited no untoward behavior during or after injections. Small intestinal lengthening was significantly greater at +1 day, +50 days and as long as 6 months after the final injection (**Figure 1A, S1A**). We used this model to assess how tissue adaptation is sustained downstream of this epithelial – innate immune cell circuit.

**Figure 1.**
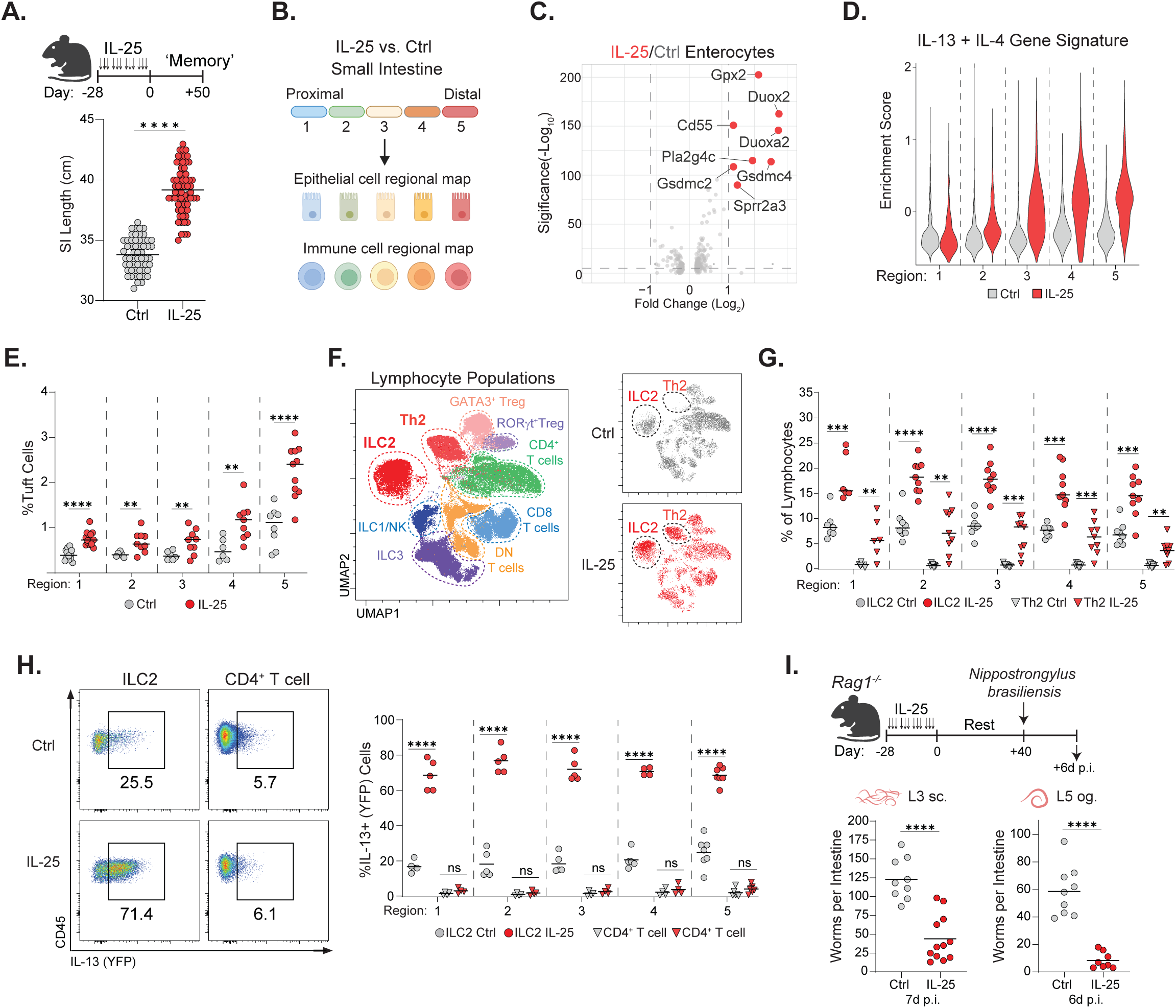
An IL-25 regimen induces long term adaptation of epithelial and immune compartments in the small intestine. (A) The IL-25 treatment regimen and small intestinal length +50 days post treatment. (B) Dividing the small intestine into 5 equal lengths for epithelial and immune cell examination. (C) Volcano plot of differentially expressed genes in enterocytes. (D) Violin plot showing the IL-13 + IL-4 gene signature enrichment score in regional enterocytes. (E) Quantification of tuft cell abundance in the small intestine. (F) Representative UMAP generated by flow cytometry of lymphocyte populations in the small intestine. (G) Quantification of ILC2 and Th2 abundance in the small intestine. (H) Representative plots and quantification of IL-13 (YFP) expression by ILC2s and CD4+ T cells in the small intestine. (I) The IL-25 treatment regimen in *Rag1^-/-^* mice followed by *N. brasiliensis* infection and intestinal worms on indicated days. Data pooled from at least two independent experiments (A, E, G, H, I) or from one experiment representative of at least 3 experiments (F, H). Each dot represents an individual mouse. Unpaired t test was performed. Statistical significance is indicated by **p < 0.01, ***p < 0.001, ****p < 0.0001; ns, not significant. See also Figure S1.

The small intestine is regionalized into five cellular and metabolic domains extending from proximal duodenum to the distal ileum in mouse and human (18). To ensure all domains were sampled, we divided the tissue into 5 equal lengths and isolated epithelial (EpCAM+CD45-) and immune cells for analysis from control or IL-25-treated mice at the +50 day time point (**Figure 1B**). Each of the epithelial cell types, including enterocytes, crypt cells, Paneth cells, goblet cells, tuft cells, and enteroendocrine cells, were present in both conditions: no unexpected populations were present and regional populations remained roughly proportional (**Figures S1B, S1C, and S1D**). In comparing differential gene expression in absorptive enterocytes under the two conditions, we found upregulation of a suite of genes induced by IL-13, IL-4 and/or helminth infection, including *Gsdmc2*, *Gsdmc4*, *Pla2g4c*, and *Sprr2a3* (**Figure 1C**) (19–21). We validated expression of Gsdmc2/3, which was abundantly expressed in enterocytes from IL-25-treated animals as compared to controls in agreement with prior studies (**Figure S1E**)(21). We grouped expression of all of these target genes into a composite IL-13/IL-4 gene signature and documented a proximal-to-distal gene gradient that was enriched among enterocytes from IL-25-treated mice (**Figure 1D**) and correlated with the increased numbers of tuft cells along a similar gradient as previously noted (**Figure 1E, S1F**)(11,12).

We next examined intestinal immune cells in the small intestine lamina propria +50 days after completing treatment with IL-25 using standard flow cytometric gating analysis (**Figure 1B, S1G**). As assessed by multiparameter spectral flow cytometry, ILC2s and Th2 cells increased in all regions of the small intestine in IL-25 treated mice; other lymphocyte populations were less affected (**Figure 1F, 1G and S1H**). Intestinal ILC2s from IL-25-treated animals displayed strong down regulation of CD25 and Sca-1, decreases in CD127 and Thy1.2, and diminished proliferation as assessed by Ki67 (**Figure S1I**). Minor phenotypic differences were present in Th2 cells from IL-25-treated animals although few Th2 cells were present in control animals (**Figure 1F,1G**). We used IL-13 and IL-5 cytokine reporter mice as previously described (11,14) to compare the functional state of ILC2s and Th2 cells from +50 day IL-25-treated and control mice. Unexpectedly, ILC2s from treated mice had sustained IL-13 reporter expression that was not present in control ILC2s or CD4^+^ T cells under either condition (**Figure 1H**); a similar trend was present for expression of the IL-5 reporter, which is expressed more highly among steady state ILC2s (**Figure S1J**)(22). Taken together, IL-25 alone induces lasting small intestinal adaptations that overlap effects seen after helminth infection, including intestinal lengthening, evidence for IL-13-induced enterocyte gene expression and increases in tuft cells and activated ILC2s.

### Group 2 ILCs are sufficient to induce lasting small intestinal adaptation

The presence of increased numbers of activated ILC2s suggested their role in small intestinal adaptation. Indeed, as assessed at +1 day, *Rag1*-deficient mice had intestinal lengthening when treated with IL-25 that was not present in lymphocyte-deficient *Rag2 IL2rg*-deficient mice (**Figure S1K**). Mice lacking IL-4 and IL-13 or the shared receptor subunit, IL4ra, fail to induce the tuft cell-ILC2 circuit in response to helminths (11) and fail to develop small intestinal lengthening in response to IL-25 (**Figure S1K**). In contrast, IL-5-deficient (*R5^Cre/Cre^* homozygous) mice lacking most eosinophils developed intestinal lengthening but this was absent when these mice were crossed with *Il17rb^ff^* mice that delete the IL-25-binding component of the IL-25 receptor from IL-5-expressing cells, which are predominantly ILC2s (**Figure S1K**). Small intestinal lengthening was sustained in IL-25-treated Rag1-deficient mice at the day +50 time point (**Figure S1L**). Taken together, we conclude that ILC2s activated to produce IL-13 by IL-25 can induce persistent small intestinal adaptation.

A cardinal feature of immune memory is increased protection against future infections. To test this, we compared matched cohorts of *Rag1*-deficient mice treated with IL-25 or control and maintained for +40 days before subcutaneous infection with migratory L3 *N. brasiliensis* larvae (**Figure 1I**). Although control mice were unable to reject worms, as occurs in the absence of adaptive lymphocytes, IL-25-treated mice had significantly fewer parasites. Additional cohorts were challenged using L5 *N. brasiliensis* larvae that had matured for intestinal life before being deposited directly into the gut by oral gavage. Again, significant protection was present in IL-25-treated mice and consistent with the presence of functionally relevant innate immune-mediated memory in the small intestine.

### IL-25 drives accumulation of activated ILC2s in additional tissues

Activation of small intestine ILC2s by helminth infection or acute injections of IL-25 drives activation, proliferation and egress into the circulation and accumulation in other organs, including the lung and adipose tissues (23,24). To assess the impact of IL-25-induced memory on ILC2s in distant organs, we compared the phenotypes of ILC2s taken from lung, adipose tissues and colon at +50 days after IL-25 or control regimens (**Figure S2A**). ILC2s were phenotypically altered in each of these tissues and more abundant in lung and colon (**Figure 2A, 2B**). Tissue ILC2s shared downregulation of CD25 and Sca-1 and similar levels of CD127 and Ki67, whereas markers like KLRG1, Thy1.2 and PD-1 varied by tissue (**Figure 2C, S2B**). ILC2s in lungs and fat of IL-25-treated mice upregulated α4β7 integrin, which binds its ligands MAdCAM-1 and VCAM-1 to promote mucosal and gut homing, although the integrin was not detected on the surface of intestinal ILC2s, possibly due to downregulation by the abundance of ligand or technical issues related to isolation of gut cells. We also used cytokine reporter mice to compare the activation state of tissue ILC2s induced by IL-25 treatment. Like small intestine conditioned ILC2s, greater proportions of ILC2s expressed IL-13 and IL-5 as compared to control mice (**Figure 2D, S2D**) while CD4+ T cell cytokine expression was unaltered (**Figure S2C, S2D**).

**Figure 2.**
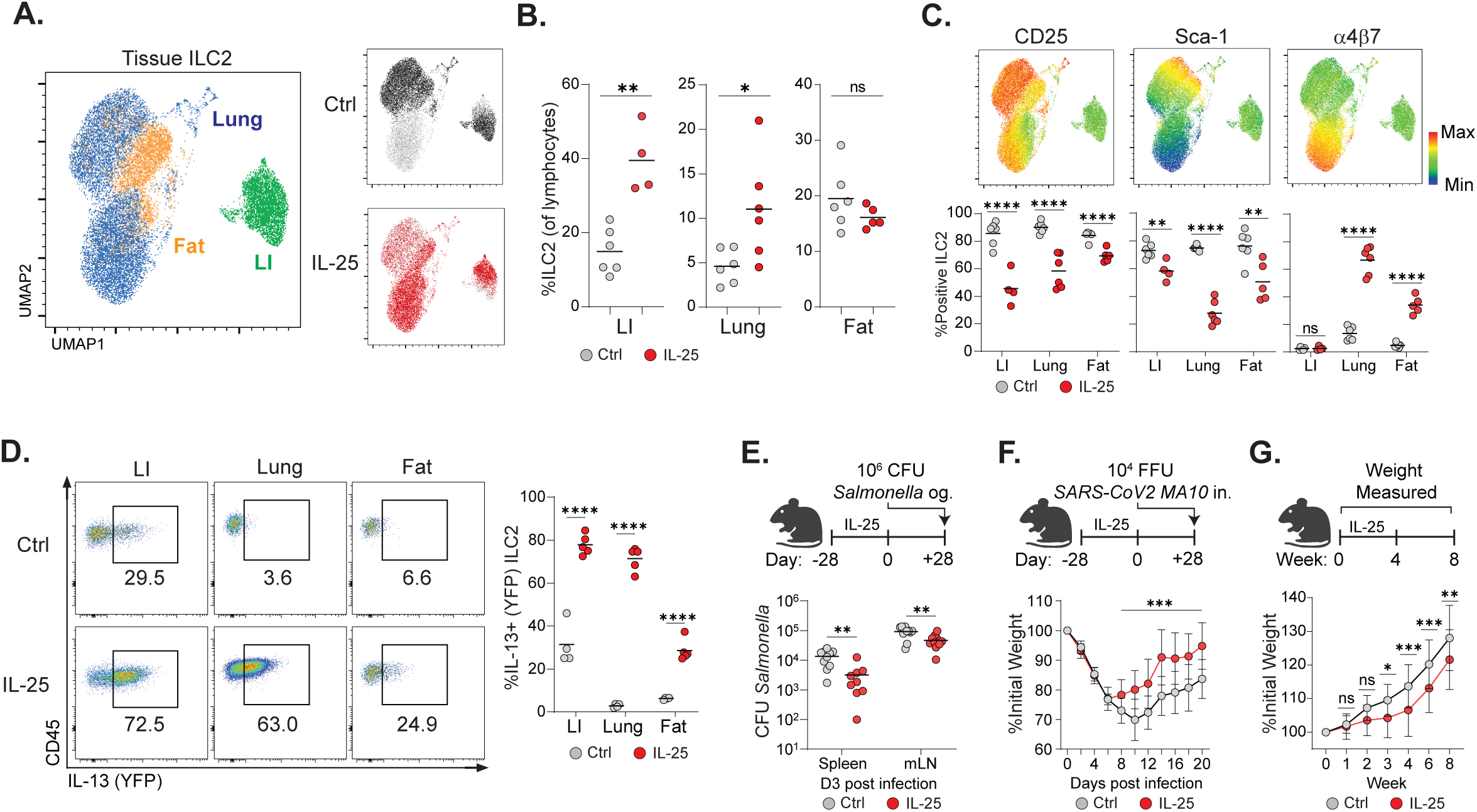
The IL-25 regimen alters ILC2 populations across tissues, enhances mucosal resilience and reduces adiposity. (A) UMAP generated by flow cytometry of ILC2s, displaying phenotype based on tissue (large intestine-LI, lung, or fat) and treatment condition. (B) ILC2 abundance in tissues from control or IL-25 treated mice. (C) Expression and quantification of indicated markers by tissue ILC2s. (D) Representative plots and quantification of IL-13 (YFP) expression by tissue ILC2s. (E) Titers of *Salmonella* in the spleen and mesenteric lymph nodes (mLN) 3 days after infection in control or IL-25 treated mice. (F) Weight loss during infection with SARS-CoV-2 MA10 in control or IL-25 treated mice. (G) Weight gain during the IL-25 regimen and 4 weeks post treatment. Data pooled from at least two independent experiments (B, C, D, E, F, G) or from one experiment representative of 3 experiments (A). Each dot represents an individual mouse (B, C, D, E); Dot represents mean ± SD, n = 15-21 (F, G). Unpaired t test (B, C, D), two-tailed t tests of area under the curve (F), or two-way ANOVA (Bonferroni’s post-hoc test) (G) was performed. Statistical significance is indicated by *p < 0.05, **p < 0.01, ***p < 0.001, ****p < 0.0001; ns, not significant. See also Figure S2.

### IL-25 conditioning enhances resilience to mucosal pathogens and reduces adiposity

The dissemination of activated ILC2s to distant tissues after helminth infection can promote mucosal protection at distant sites, including to unrelated helminths (9). To assess whether treatment with IL-25 alone could establish protection, we examined the response of mice to an enteric bacterial pathogen, *Salmonella typhimurium* SL1344. Mice were induced with IL-25, rested for 28 days and infected with *Salmonella* by oral gavage (**Figure 2E**). As assessed 3 days after bacterial challenge, IL-25 induced animals had significantly reduced numbers of bacteria in spleen and mesenteric lymph nodes (mLN) as compared to control animals. Recently, prior lung-migratory helminth infection was shown to improve outcomes of SARS-CoV2 lung infection (25). When challenged using mouse-adapted SARS-CoV2 MA10 virus at day +28 after induction, mice conditioned with IL-25 had significantly earlier recovery from weight loss as compared to control animals although improved survival did not reach significance (**Figure 2F, S2E**). Finally, the presence of activated ILC2s in adipose tissue suggested metabolic adaptations as previously described in IL-33-treated animals (26,27). Indeed, IL-25-treated mice had reduced weight gain which persisted up to four weeks after treatment in association with reduced adipose tissue and preservation of lean body mass (**Figure 2G, S2F**). Finally, we tested whether the presence of increased number of activated ILC2s in small intestine would render animals more susceptible to type 2 pathology. To test this, we sensitized IL-25-conditioned and control mice after +45 days with oral ovalbumin (OVA) and cholera toxin to establish allergic sensitization via the gastrointestinal mucosa (**Figure S2G**)(28). Systemic challenge with OVA at +60 days elicited comparable falls in core body temperature and increases in serum mast cell MCTP-1 in the two cohorts, suggesting little change in allergic sensitization after IL-25 adaptation as assessed in small intestine.

Taken together, small intestinal adaptation elicited by IL-25 alone can induce mucosal and metabolic changes reported after viable helminth infection while avoiding pathology due to the latter. Of note, however, acute mucosal co-infections with helminths and heterologous pathogens can result in worse outcomes due to impairment of appropriate host immune responses (29). To assess whether acute treatment with IL-25 alone creates comparable windows of vulnerability, we treated cohorts of mice with 3 injections of IL-25 over 1 wk followed by one day of rest (acute), or 12 injections of IL-25 over 4 wks followed by one day of rest (adapted), before the indicated challenges. After infection with *Salmonella*, acute mice had significantly more bacteria in spleen and mLNs as compared to adapted mice (**Figure S2H**). Similarly, adapted mice challenged with SARS-CoV-2 had less weight loss and increased survival as compared to challenged acute mice (**Figure 2SI**). We conclude that mucosal and metabolic alterations initiated by IL-25 proceed through a window of pathogen vulnerability as tissues undergo the cellular and morphologic changes that stabilize the adapted state.

### IL-25 conditioned ILC2s have a distinct transcriptional profile

To characterize the transcriptional profile of ILC2s at the memory time point (+50 days), we performed scRNA-seq on small intestinal ILC2s purified from IL-25 treated or control mice (**Figure S3A**). As visualized by reduction and clustering in UMAP space, control and IL-25 induced ILC2s expressed divergent transcriptomic signatures, with differentially expressed genes represented as a volcano plot (**Figure 3A, 3B**). Type 2 cytokines like IL-13, IL-4 and IL-5 were increased in IL-25 conditioned ILC2s as supported by prior analysis using cytokine reporter mice (**Figure 3C**). GM-CSF in the adjacent locus, however, was attenuated and validated using cells from *Csf2* reporter mice (**Figure S3B**); amphiregulin and IL-9 were little different.

**Figure 3.**
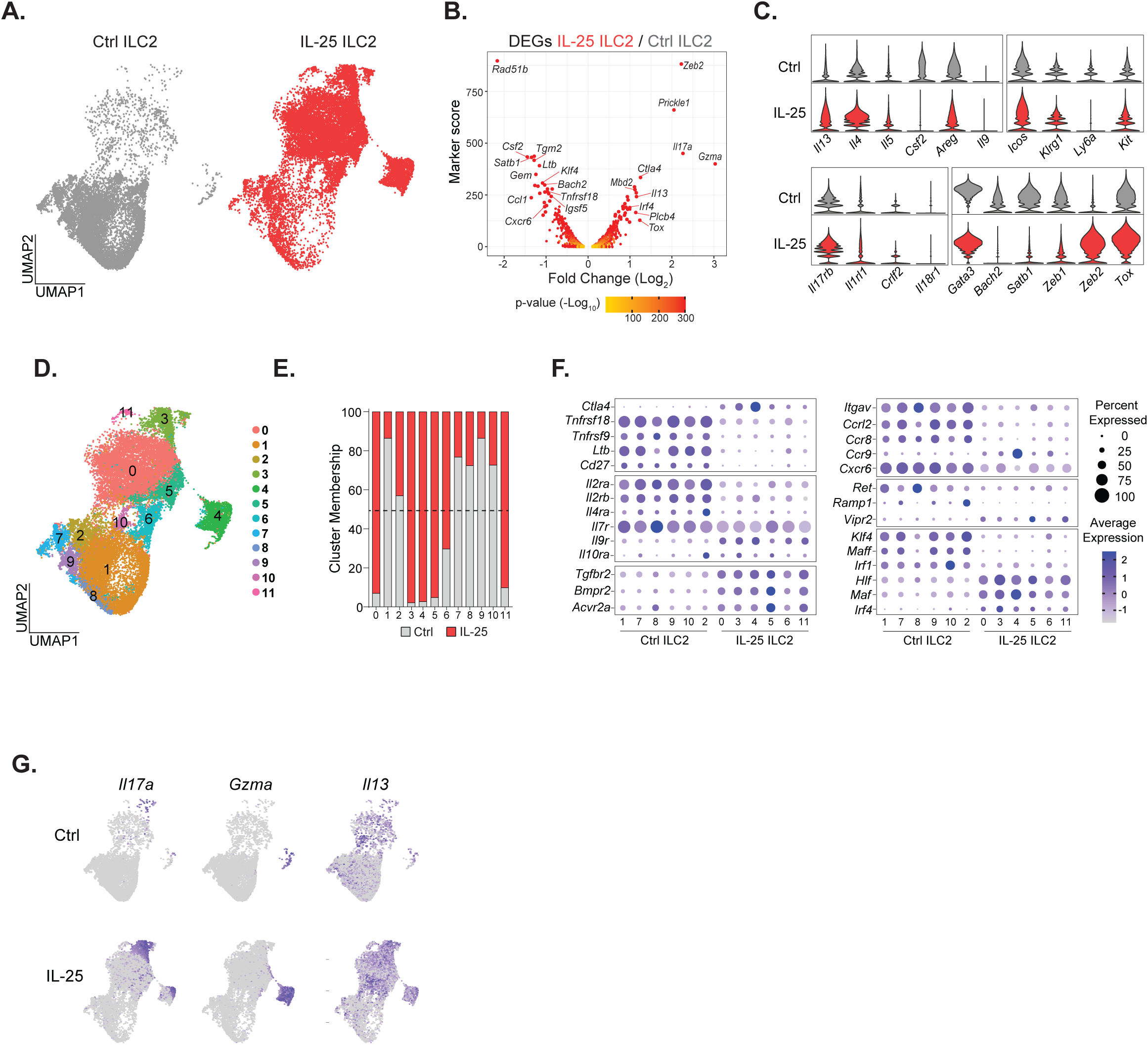
ILC2s maintain a distinct transcriptional state after IL-25 induction. (A) UMAP generated by scRNA-seq of small intestinal ILC2s at +50 d after control or IL-25 treatment. (B) Volcano plot showing differentially expressed genes between ILC2s collected under indicated conditions. (C) Violin plots showing expression of indicated genes in ILC2s. (D) UMAP generated by scRNA-seq showing ILC2 populations. (E) ILC2 cluster representation in control or IL-25 treatment conditions. (F) Dot plots showing expression of indicated genes by ILC2 populations. (G) Feature plots showing expression of indicated genes on the ILC2 UMAP space. See also Figure S3.

Transcripts for surface markers such as *Icos*, *Klrg1*, and *Kit* were similarly expressed, whereas *Ly6a* (which encodes Sca-1) was downregulated, as verified by flow cytometry (**Figure S1I**). ILC2s from both conditions had similar expression of the alarmin receptors *Il17rb*, *Il1rl1*, *Crlf2*, and *Il18r1* (receptors for IL-25, IL-33, TSLP, and IL-18, respectively), with *Il17rb* and *Il1rl1* being most abundant. While IL-25 conditioned ILC2s maintained high expression of the lineage defining transcription factor *Gata3*, we noted dynamic changes in other transcription factors, including reductions in *Bach2*, *Satb1*, and *Zeb1*, and marked upregulation of *Zeb2* with modest upregulation of *Tox*.

We used clustering analysis to visualize transcriptionally distinct populations of ILC2s (**Figure 3D, S3C**). This analysis generated 12 clusters, with clusters 1, 7, 8, 9, and 10 predominant in control animals, clusters 0, 3, 4, 5, 6, and 11 in IL-25 treated animals, and cluster 2 in both conditions (**Figure 3E**). Each cluster expressed some genes which were unique as compared to the other clusters (**Figure S3C**). However, many genes implicated in ILC2 function showed expression patterns shared among clusters either from control or from IL-25 treated mice with cluster 2 grouping with control clusters (**Figure 3F**). Genes of interest upregulated in IL-25 conditioned ILC2s were inhibitory receptor *Ctla4*, the cytokine receptors *Il9r* and *Il10ra*, the TGFβ family cytokine receptors *Tgfbr2*, *Bmpr2*, and *Acvr2a*, the chemokine receptor *Ccr9*, the neuropeptide receptor *Vipr2*, and the transcription factors *Hlf*, *Maf*, and *Irf4*.

Of note, ILC2 clusters 3 and 4 expressed transcripts for *Il17a* and *Gzma,* which are genes usually associated with ILC3s and ILC1s, respectively (**Figure 3G, S3C**). These *Il17a* and *Gzma* expressing ILC2s maintained expression of core ILC2 identity genes such as *Il13* and *Gata3* and did not express other genes typical of ILC3s or ILC1s, including *Rorgt*, *Tbx21*, *Eomes*, *Il22*, or *Ifng* (**Figure 3G, S3D**). Thus, IL-25 induction results in emergence of distinct populations of ILC2s in the small intestine that survive for many months and display heightened effector capacity (*Il13*, *Il4*, and *Il5*) and novel capabilities (*Il17a* and *Gzma*), which are features of immune cell memory.

### Tissue adaptation and memory ILC2s persist independent of endogenous alarmins or tuft cells

We hypothesized that small intestinal adaptation could be sustained by the feed-forward nature of the tuft cell-ILC2 circuit via IL-25 from the increased numbers of tuft cells (**Figure 1E**). To test this, we treated *Il25^-/-^* mice with IL-25 and assessed attributes of small intestinal adaptation at +50 days (**Figure 4A**). Unexpectedly, treated IL-25-deficient mice developed small intestinal lengthening, increased numbers of tuft cells and a greater abundance of ILC2s (**Figure 4A, S4A**). Additionally, we performed scRNA-seq from ILC2s collected across small intestine from WT and *Il25^-/-^* mice after induction with control or IL-25 at +50 days (**Figure S4B, 4B**), and revealing populations of IL-25-dependent ILC2s that are missing in *Il25^-/-^* mice under resting conditions (**Figure 3A, 4B, S4C, S4D**; WT vs IL-25-/-control mice). After IL-25 induction, WT and *Il25^-/-^* ILC2s occupy essentially identical transcriptional space including increased expression of *Il13*, *Il4*, and *Zeb2*, as well as reduced expression of *Csf2*, *Rad51b*, and *Sat1b* (**Figure 4C**). We documented the accumulation of specialized subsets that expressed transcripts for *Il17a*, *Gzma* and, more generally, *Il13* (**Figure S4E**), the differential expression of CD25, Sca-1, KLRG1 and PD-1 using spectral flow cytometry (**Figure S4F**) and confirmed the activated ILC2 phenotype as assessed using mice crossed to reporter alleles for IL-13 and IL-5 (**Figure 4D, S4G**). Since intestinal tuft cells can respond to alternative parasite ligands and produce additional factors besides IL-25 that activate ILC2s (30,31), we administrated IL-25 to *Pou2f3^-/-^* mice, which lack tuft cells. Again, small intestinal lengthening and increased proportions of ILC2s occurred at +50 days despite the absence of tuft cells (**Figure S4H, S4I**). IL-25 conditioned ILC2s from *Pou2f3^-/-^*mice also shared phenotypic alterations seen in wild type and *Il25^-/-^*mice (**Figure S4J**). Additionally, ILC2s from *Pou2f3*^-/-^ mice had increased expression of IL-13 and IL-5 consistent with persisting activation (**Figure S4K**). We conclude that IL-25 acting via ILC2s is required to induce small intestinal adaptation (**Figure S1K**) but that neither IL-25 nor tuft cells is required to maintain it.

**Figure 4.**
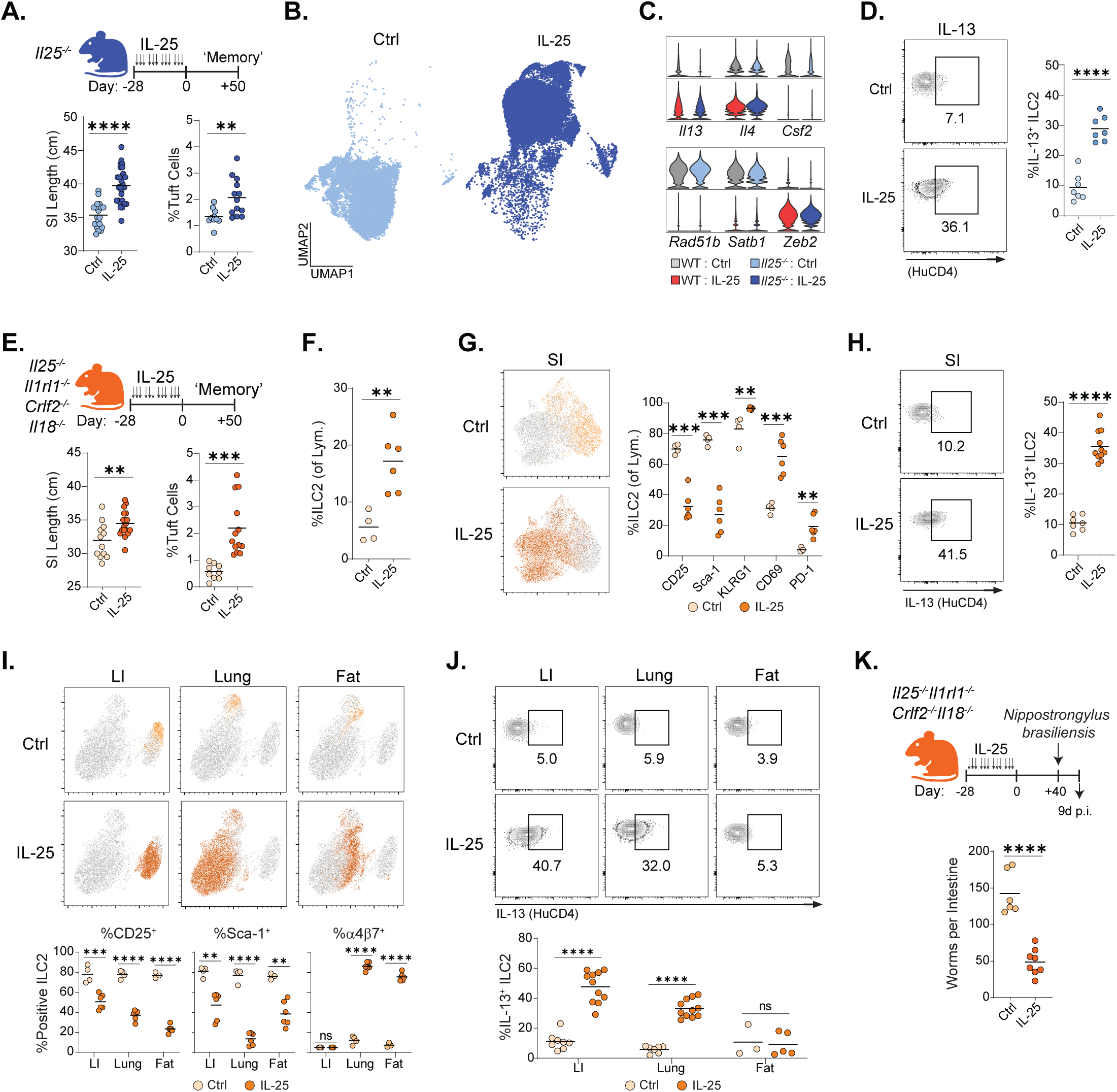
Small intestinal adaptation is maintained independently from endogenous alarmin cytokines. (A) The IL-25 treatment regimen in *Il25^-/-^* mice and quantification of small intestinal length and tuft cell abundance. (B) UMAP generated by scRNA-seq of ILC2s from *Il25^-/-^* mice after the IL-25 regimen. (C) Violin plots showing expression of indicated genes in wild type or *Il25^-/-^* ILC2s. (D) Expression of IL-13 (HuCD4) by small intestine ILC2s from *Il25^-/-^* reporter mice. (E) The IL-25 treatment regimen in *Il25^-/-^Il1rl1^-/-^Crlf2^-/-^Il18^-/-^* mice and quantification of small intestinal length and tuft cell abundance. (F) ILC2 abundance in the small intestine of *Il25^-/-^Il1rl1^-/-^Crlf2^-/-^Il18^-/-^* mice. (G) UMAP generated by flow cytometry of ILC2s from the small intestine of *Il25^-/-^Il1rl1^-/-^Crlf2^-/-^ Il18^-/-^* mice. (H) Expression of IL-13 (HuCD4) by small intestine ILC2s from *Il25^-/-^Il1rl1^-/-^Crlf2^-/-^Il18^-/-^* mice. (I) UMAP generated by flow cytometry of ILC2s from the large intestine (LI), lung, and fat of *Il25^-/-^Il1rl1^-/-^Crlf2^-/-^Il18^-/-^* mice after the IL-25 regimen and quantification of indicated markers. (J) Expression of IL-13 (HuCD4) by LI, lung, and fat ILC2s from *Il25^-/-^Il1rl1^-/-^Crlf2^-/-^Il18^-/-^* mice. (K) The IL-25 treatment regimen in *Il25^-/-^Il1rl1^-/-^Crlf2^-/-^Il18^-/-^* mice followed by *N. brasiliensis* infection. Intestinal worms 9 d after infection. Data pooled from at least two independent experiments (A, D, E, F, G, H, I, J, K). Each dot represents an individual mouse (A, D, E, F, G, H, I, J, K). Unpaired t test was performed. Statistical significance is indicated by **p < 0.01, ***p < 0.001, ****p < 0.0001; ns, not significant. See also Figure S4.

Although IL-25 is a key upstream activating cytokine for small intestinal ILC2s, the induced ILC2s continued to express receptors for multiple alarmins implicated in establishing thresholds for type 2 cytokine expression in tissue ILC2s and Th2s (**Figure 3C**)(32). Further, gasdermin C, which was highly expressed, helps orchestrate the alarmin response to intestinal helminth infection (21). To assess roles for alarmins in the activated tissue phenotype of the IL-25-induced ILC2s, we generated mice deficient in IL-25, ST2 (the IL-33 receptor component, *Il1rl1*), TSLPR (TSLP receptor component, *Crlf2*) and the cytokine IL-18 that were crossed further to IL-13 and IL-5 cytokine reporters to assess the activation state of tissue ILC2s *in situ*. Unexpectedly, *Il25^-/-^ Il1rl1^-/-^ Crlf2^-/-^ Il18^-/-^* mice treated with IL-25 and assessed at least 50 days later developed anatomic (lengthening), cellular (increased ILC2s and tuft cells), and ILC2 phenotypic (decreases in CD25 and Sca-1 with increases in CD69 and PD-1) and activation markers (increased IL-13 and IL-5) associated with WT ILC2s previously induced with IL-25 (**Figure 4E, 4F, 4G, 4H, S4L**). Thus, IL-25 induced ILC2s are tissue-resident cells sustained in an activated state independent of alarmins associated with type 2 allergic pathology.

ILC2s in tissues like the lung and adipose constitutively express ST2 and are more dependent on IL-33 for their basal activation than small intestinal ILC2s (23,24). Despite this, ILC2s from large intestine, lung and gonadal adipose tissue of IL-25-conditioned quadruple alarmin-deficient mice acquired the same phenotype as ILC2s from WT mice induced with IL-25 and examined 50 or more days later. ILC2s increased in lung and colon (**Figure S4M**) and were phenotypically distinct from ILC2s in control mice as assessed by expression of CD25, Sca-1, KLRG1, CD69 and PD-1 (**Figure 4I, S4N**), as well as elevated integrin α4β7 in conditioned ILC2s from lung and adipose tissue (**Figure 2C, 4I**). Further, IL-25-conditioned ILC2s from *Il25^-/-^ Il1rl1^-/-^ Crlf2^-/-^ Il18^-/-^* mice showed evidence for activation as assessed by increased IL-13 reporter expression in lung and colon and IL-5 expression in these sites as well as in adipose tissue (**Figure 4J, S4O**). Finally, we assessed the functional relevance of IL-25-induced memory in alarmin-deficient animals using challenge with *N. brasiliensis* 40 days after the final dose of IL-25 or control (**Figure 4K**). Although *N. brasiliensis* control is almost completely alarmin-dependent in WT mice (33), IL-25 conditioned mice lacking alarmins acquired substantial immunity against helminth challenge as assessed 9 days after challenge that was absent in control animals. We conclude that IL-25 conditioned mice develop robust mucosal adaptation that is sustained independently from the alarmin inputs required for primary responses to helminth challenge.

### IL-25 conditioned ILC2s display sustained epigenetic alterations

The sustained phenotypic and transcriptional changes in ILC2s conditioned by prior IL-25 administration suggested that epigenetic adaptations may be important in maintenance of the ILC2 population. To assess this, we performed bulk ATAC sequencing on small intestinal ILC2s at +50 days, revealing over 15,000 differentially expressed peaks as compared to ILC2s from control mice isolated and examined concurrently (**Figure 5A, S5A**). Genetic accessibility was altered across the genome predominantly within intergenic, promoter and intronic regions (**Figure 5B, 5C**). The type 2 cytokine region of chromosome 11 revealed increased accessibility of regions for *Kif3a*, *Il4*, *Rad50*, the 3’ region of *Il13*, and distal 5’ region of *Il5* although excluding *Csf2*, as suggested by reduced transcript levels recovered in the scRNAseq analysis (**Figure 5D, 5E, 3C**). Although some of these align with well-described regulatory sequences (34), several were in previously unrecognized genomic regulatory regions, although these continue to be experimentally defined (35). Accessibility of the *Il17a* and the *Gzma* loci was consistent with the transcript data (**Figure 5F, 3G**), although we noted no increase in accessibility of transcription factors *Rorc* and *Tbx21*, which can drive IL-17a and granzyme expression, respectively (**Figure S5B**). Taken together, IL-25 conditioned ILC2s acquire an epigenetically differentiated state across genomic loci of relevant immune function and consistent with a state of memory.

**Figure 5.**
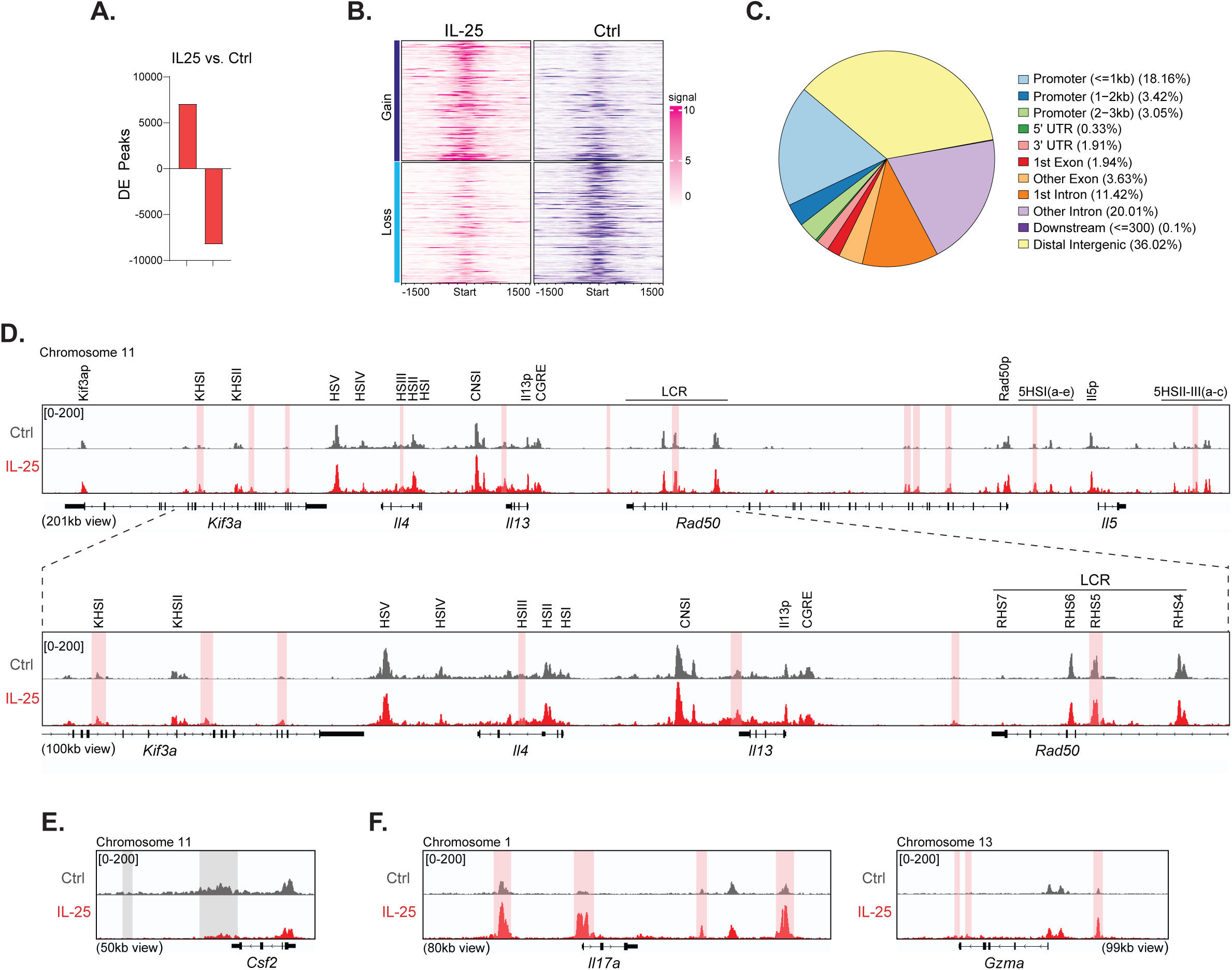
IL-25 conditioned ILC2s display epigenetic hallmarks of memory. (A) The number of differentially expressed peaks between control or IL-25 conditioned ILC2s from the small intestine. (B) Plot showing the gain and loss of peaks across the genome in ILC2s. (C) Pie chart showing genomic regions of differentially expressed peaks. (D) Genomic tracts of selected areas within the type 2 cytokine locus. Indicated Regulatory Sequences from references 34,35. Shaded areas represent regions with significantly more peaks (pink) or less peaks (grey) in IL-25 conditioned ILC2s. (E) Genomic tracts of the *Csf2* locus. (F) Genomic tracts of the *Il17a* locus and the *Gzma* locus. See also Figure S5.

### Memory ILC2s sustain intestinal barrier alterations through IL-4ra signaling

Whether tissue adaptation is sustained collectively by multiple cell types or selectively by specific cell types remains an active area of investigation that likely depends on the nature of the inflammation and tissue(s) involved (36). Epithelial stem cells have been implicated as central mediators sustaining tissue adaptation in several types of inflammation, either directly or with input from the microbiota (4,5,37); glial astrocytes, barrier cells of the central nervous system, may subserve a similar role (38). To address contributions by the microbiota, we co-housed IL-25-and control-treated cytokine reporter mice throughout induction and follow-up but noted no impact on small intestinal lengthening or accumulation of activated ILC2s in IL-25-treated mice (**Figure S6A**). Reconstitution of germfree mice using cecal content from IL-25-treated mice had no impact on intestinal lengthening, tuft cell abundance or ILC2 numbers (**Figure S6B**), and IL-25 was sufficient to induce small intestinal lengthening in germfree mice (14). *Tritrichomonas* cecal colonization increased basal tuft cell and ILC2 numbers but IL-25 induction resulted in the same final increases in intestinal length, tuft cell and ILC2 accumulation and activation as seen in treatment of *Tritrichomonas*-free animals, as previously noted (14). Thus, contributions from the microbiota, at least under these conditions, are alone unlikely to account for the adaptation that occurs following IL-25 administration.

To assess a role for epithelial stem cells in sustaining IL-25-conditioned adaptation, we isolated intestinal stem cells from *Lgr5*-EGFP reporter mice across the small intestine at the +50 day time point for analysis by ATAC sequencing (**Figure S6C**). This population of epithelial cells from control and IL-25 adapted mice had five differentially expressed peaks and were essentially indistinguishable (**Figure S6D**). To assess the status of crypt epithelial transit amplifying cells in response to IL-25 (39,40), we refined clustering of crypt-derived cells in the scRNA-seq epithelial data set to resolve 4 populations that could be separated into *Lgr5*-high and populations driven by differential expression of *Olfm4*, *Mki67*, and *Cd44* (**Figure S1B, S6E**). When each population was analyzed, *Lgr5*^+^ cells showed minimal induction of the IL-13 gene signature that was induced to variable amounts in the three other crypt-associated populations (**Figure S6F, 1D**). These findings are consistent with a role for exogenous IL-13/IL-4 in driving the alterations of epithelial gene expression and cellular composition during epithelial differentiation among transit-amplifying cells but not crypt stem cells.

To further assess this, we generated organoids from crypt cells isolated from proximal, mid and distal small intestine from control and IL-25-conditioned mice and used flow cytometry to examine tuft cell frequency in the absence or presence of exogenous IL-13 to assess sensitivity to stimulation by type 2 cytokines. Under these conditions, there was no difference in tuft cell abundance in organoids derived from control or IL-25-conditioned mice in the absence or presence of IL-13 (**Figure 6A**). We co-cultured crypt-derived organoids from control or IL-25-conditioned mice with small intestinal ILC2s isolated from control or IL-25-conditioned mice and examined the abundance of tuft cells 6 days later. Regardless of the origin of crypt cells, increases in tuft cell abundance depended on co-culture with IL-25-conditioned memory ILC2s (**Figure 6B**). We verified that the numbers of ILC2s was similar at the end of the culture period among each of the groups (**Figure S6G**), suggesting that the activation state of the IL-25 conditioned ILC2s was responsible for the phenotype. Indeed, addition of IL-13 and IL-4 neutralizing antibodies to the organoid-ILC2 co-cultures reduced the numbers of tuft cells while control antibodies had no effect (**Figure 6C**). Thus, as assessed in this *in vitro* system, intestinal epithelial adaptation is sustained by memory ILC2s and not by alterations of the epithelial stem cell compartment.

**Figure 6.**
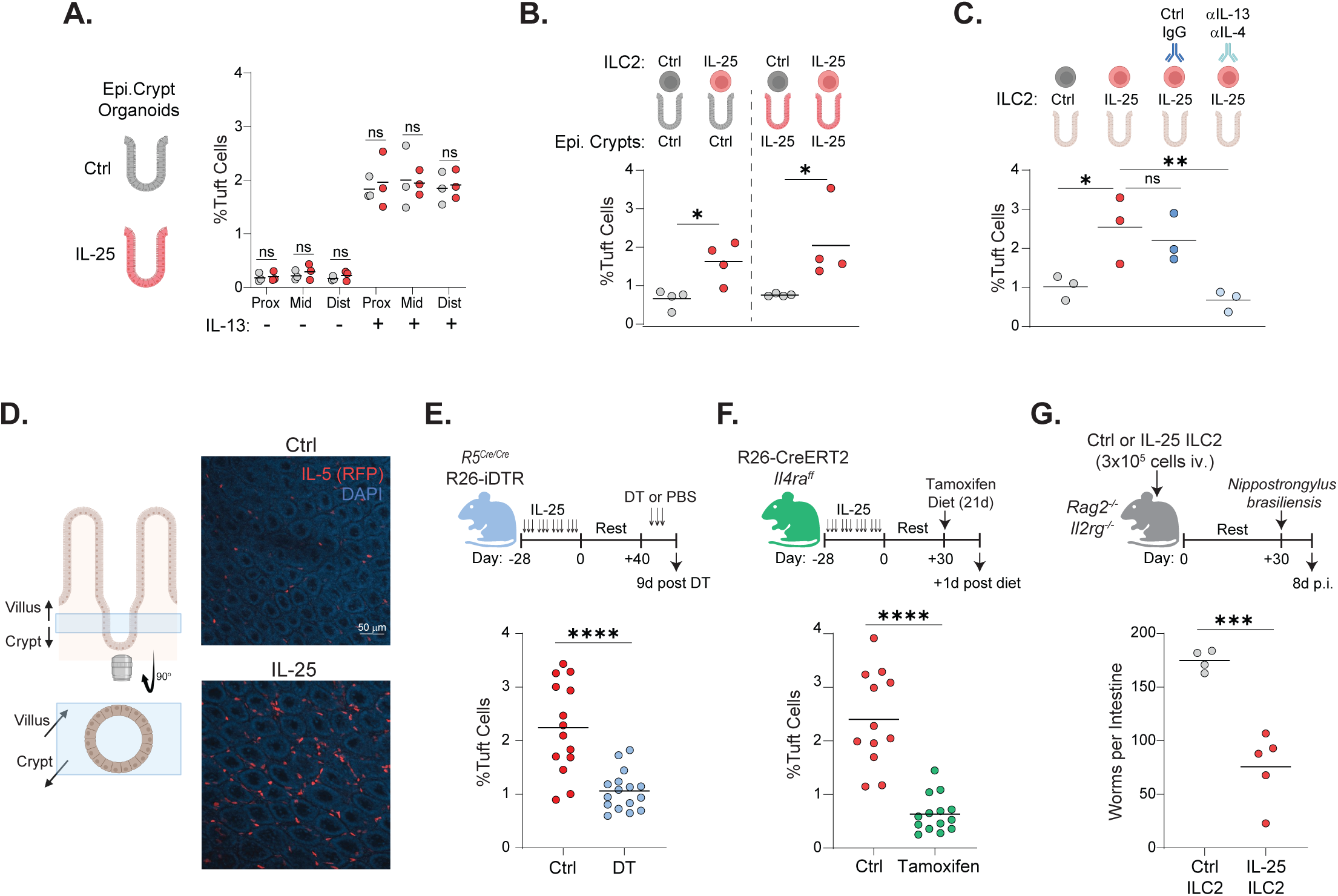
Memory ILC2s sustain small intestinal adaptation. (A) Organoids of epithelial (Epi.) crypts from the proximal, mid, and distal small intestine of mice +50 d after control or IL-25 treatment. Tuft cell abundance was quantified by flow cytometry under standard organoid culture with or without added IL-13. (B) Organoids of control or IL-25 conditioned ILC2s with Epi. crypts from control or IL-25 treated mice. Tuft cell abundance was quantified. (C) Culture of control or IL-25 conditioned ILC2s with passaged crypt organoids. Control IgG or anti-IL-13 and anti-IL-4 was added to cultures with IL-25 conditioned ILC2s. Tuft cell abundance was quantified. (D). Representative image of IL-5-expressing cells localized within the crypt-villus region (blue box) of the distal small intestine of control or IL-25 treated mice. (E) Abundance of tuft cells in small intestine of R5-Cre-R26-iDTR mice after ILC2 depletion. (F) Abundance of tuft cells in small intestine of R26-CreERT2 x *Il4ra^ff^* mice after IL-4ra deletion. (G) Transfer of ILC2s to *Rag2^-/-^ Il2rg^-/^*^-^ mice followed by *N. brasiliensis* infection. Intestinal worms 8 days after infection. Data pooled from at least two independent experiments (A, B, C, E, F, G), or from one representative experiment (D). Each dot represents an individual mouse (A, B, C, E, F, G); Each dot is the average of 3 technical replicates (A, B, C). Unpaired t test (A, B, E, F, G), or one-way ANOVA (Dunnett’s post-hoc test)(C) was performed. Statistical significance is indicated by *p < 0.05, **p < 0.01, ****p < 0.0001; ns, not significant. See also Figure S6.

As assessed by immunohistochemistry, memory ILC2s expressing IL-5 were expanded in areas adjacent to the transit amplifying zone at the crypt-villus axis but also dispersed along the length of the villus where effector cytokines could impact gene expression in adjacent epithelial cells (**Figure 6D, S6H**). To assess whether memory ILC2s sustain the epithelial phenotype *in vivo*, we used R5^Cre/Cre^ x R26-iDTR mice to drive diphtheria toxin receptor (DTR) expression on IL-5-expressing cells. After IL-25 induction and rest, mice were treated 3 times with diphtheria toxin (DT), which caused almost complete loss of small intestinal ILC2s (**Figure S6I**) and a significant reduction in tuft cell frequency as compared to controls (**Figure 6E**). We performed similar experiments using R26-CreERT2 x Il4ra^ff^ mice, in which IL-4ra is deleted when cells are exposed to tamoxifen (**Figure 6F**). At +30 days after completing induction, mice were placed on tamoxifen-containing chow for 3 weeks. As controls, a cohort of R26-CreERT2 x IL4ra^ff^ mice was kept on standard diet or Il4ra^ff^ mice (with no CreERT2) were treated, rested, and put on tamoxifen diet. Deletion of IL4ra resulted in loss of tuft cell abundance in the IL-25-conditioned bowel, and consistent with a requirement for activated memory ILC2s to sustain the adapted small intestinal epithelial barrier.

To assess the stability of memory ILC2s *in vivo*, we purified small intestinal ILC2s from IL-25 conditioned and control mice and transferred 3 x 10^5^ cells into *Rag2 IL2rg*-deficient mice. After 30 days, mice were infected with *N. brasiliensis* and examined 8 days later (**Figure 6G**). Mice which received memory ILC2s had a significantly reduced intestinal worm burden compared to mice which received control ILC2s. Small intestinal ILC2 numbers and proliferation were similar in both groups, and the lower expression of Thy1.2 on IL-25-induced ILC2s was sustained (**Figure S6J**). Numbers of small intestinal tuft cells and small intestinal lengthening were not significantly altered (**Figure S6K**) although this may reflect the relatively small numbers of cells and the timing of analysis after helminth challenge.

## Discussion

Immunologic memory, classically associated with accumulation of adaptive, antigen-specific T and B cells, includes myeloid and innate lymphoid cells that acquire stable transcriptional, epigenetic and metabolic adaptations that increase responses to secondary challenge (3,41). More recently, studies of barrier tissue perturbations in mice and humans have characterized intrinsic epigenetic and transcriptional changes among epithelial stem cells that enable more rapid repair following injury but also increase the potential to contribute to inflammatory states (4,5,37). In an example of type 2 memory, polyp basal epithelial cells from allergic patients with chronic rhinosinusitis with nasal polyposis displayed impaired differentiation and evidence for extrinsically enforced activation of IL-13-induced genes, as seen in our studies, but intrinsic activation of Wnt pathway genes that was sustained in *ex vivo* epithelial cultures (5). In all of these studies, important mechanistic questions remain to understand how diverse immune and non-immune cells communicate to sustain tissue adaptation that preserves functional homeostasis of a given organ. Our studies suggest a role for ILC2s in coordinating an integrated mucosal barrier protective response across tissues exposed to the post-birth environment.

We used an evolutionarily tuned model of small intestinal homeostasis driven by ubiquitous, chronic interactions with soil-transmitted parasitic helminths that inhabit this tissue for reproduction and dispersal of offspring. We used recombinant IL-25 to separate putative protective host pathways that can be targeted by helminths (42) from potentially inflammatory responses driven by viable organisms and eggs. The cytokine IL-25, previously designated IL-17E, is a member of the evolutionarily ancient IL-17 family that appeared in tunicates over 500 million years ago (43). Family members are ubiquitous among marine invertebrates and vertebrates, including jawed and jawless fish (44), and are induced by intestinal injury and bacterial challenge (45). IL-17-like cytokines in *Caenorhabditis elegans* mediate responses to noxious stimuli, including food cues, revealing ancient connections with intestinal sensing (46).

Memory innate lymphoid cells share tissue residency, metabolic and epigenetic adaptations (41), although memory ILC2s have been described primarily after perturbations in lung where the transcriptomic and phenotypic characterizations differ from the populations described here (47–50). Functionally, prior characterized memory ILC2s mediated increased type 2 inflammatory outputs and enhanced allergic immunity and/or fibrosis *in vivo*. Prior models used induction with complex allergens, fungi, papain and house dust mite antigens (HDM), and typically shared a requirement for IL-33. IL-33 alone provoked functional memory lung ILC2s in association with up-regulation of the IL-25 receptor (47) in a response later linked with induction of innate training by c-Maf (49). The populations of memory ILC2s driven by IL-33 may differ from those driven by IL-25, in part shaped by the constitutive high expression of the IL-25 receptor on small intestinal ILC2s (51). We used IL-25 to avoid bioactive constituents contained in complex antigens and organisms (52), and suspect the ILC2s that accumulate represent naturally generated populations, since (1) similar cells were present in WT mice but lacking in IL-25-deficient mice, and (2) IL-25 induction resulted in identical terminal ILC2 effector populations in both WT and IL-25-deficient mice, suggesting a normal trajectory under *in situ* and cytokine-induced conditions.

In the IL-25-induced model we describe, intestinal epithelial crypt stem cells remained unaltered as revealed by ATAC-seq and by organoid cultures that did not differ between induced and control mice in either the basal or IL-13-treated state. In contrast, memory ILC2s sustained IL-13-induced effects on the barrier for many months marked by changes in cell composition and stable small intestinal lengthening while sparing crypt stem cells. Given the capacity of IL-25 and intestinal helminth infection to drive exodus of activated small intestinal ILC2s into blood and distant tissues (23,24), we speculate that ILC2s that accumulate in different organs originate from ILC2s conditioned in the small intestine, although we cannot rule out contributions from IL-25 receptor^+^ precursors from bone marrow or other tissues (53,54).

Unexpectedly, the memory ILC2s induced by IL-25 did not require IL-25 or additional canonical alarmins to sustain small intestinal adaptation or mediate enhanced anti-helminth immunity against *N. brasiliensis*, which requires alarmins for efficient control in the naive state (33). The single-cell RNA sequencing and multiparameter flow analysis in untreated, IL-25-deficient and IL-25-induced mice revealed an IL-25-dependent pathway for generating ILC2s capable of mediating IL-13/4-dependent trophic changes in small intestine that achieved independence from canonical alarmin-mediated activation signals, consistent with differentiation plasticity associated with altered epigenetic landscapes as posited over 50 years ago (55). Several different states were represented in our studies, including distinct *Il17a* and *Gzma* expressing subpopulations, consistent with prior studies (54), and further investigation will be required to establish whether each of these populations mediates the memory state or whether distinct populations participate in spatially or functionally restricted activities. We could not document increased sensitivity to food allergy despite the increased prevalence of activated ILC2s and tuft cells in small intestine. We speculate that alarmin-independence of activated intestinal ILC2s may represent the state established in natural animal populations and perhaps account for the general lack of allergic pathology in animals under native conditions as established by reduced access across the altered barrier.

Transcription factor profiling revealed not only c-Maf, previously implicated in ‘trained’ ILC2s by stabilizing the increased output of type 2 cytokines (49), but also a profile that resembled some aspects of tissue resident memory-effector T cells as characterized by reductions in Bach2 and Satb1, increased expression of TGF-β−family receptors and increases in Irf4 and Zeb2 (56). Despite expression of PD-1 and CTLA-4, IL-25-induced ILC2s are not exhausted as demonstrated by their activated cytokine profiles *in vivo*, *in vitro* and in transfer experiments. ZEB2 and HLF are transcription factors previously linked with relevant memory and effector states. Better known for a role in epithelial-mesenchymal transformation, ZEB2 (zinc finger E-box binding homeobox 2) is a transcriptional repressor (57) implicated in sustaining identity of tissue resident macrophages (58) and long-lived terminal effector/memory cell populations of CD8 T cells, iNKT and DC2 cells (59–61). HLF (hepatic leukemia factor) is a CLOCK-controlled proline-and acidic amino acid-rich basic leucine zipper (PAR bZip) factor that couples circadian metabolism of xenobiotics to tissue oscillators and optimizes lipid/cholesterol cycling and stem cell protection (62,63). Further work is needed to characterize the role of these transcription factors in IL-25-induced memory ILC2s given their persistence, effector function and potential role in metabolic pathways previously linked with tissue resilience and intestinal protection (64). Our findings demonstrate that the complexities of the host immune response can be parsed to uncover tissue protective pathways that may avoid inflammatory, metabolic and organ dysfunctional responses induced by viable pathogens.

## ACKNOWLEDGEMENTS

We thank M. Ji, Z.-E. Wang and M. Conseco for technical expertise and mouse colony maintenance; O.A. Aguilar for help creating the spectral flow antibody panels; J. Turnbaugh and the Mouse Metabolism Core at the UCSF Diabetes Research Center and Nutrition Obesity Research Center for help with metabolic experiments as supported in part by NIH P30DK098722; W. Eckalbar and the UCSF Genomics CoLab for help with scRNA-seq and ATAC-seq analysis; K.M. Ansel, A. Ma and A.B. Molofsky for pre-review of the manuscript; and members of the Locksley lab for discussion. Figures diagrams were created using BioRender. This work was supported by a Cancer Research Institute Irvington Postdoctoral Fellowship (V.S.C.), grants from the National Institutes of Health (AI026918, HL107202 to R.M.L.) and the Howard Hughes Medical Institute (R.M.L.).

## AUTHOR CONTRIBUTIONS

V.S.C conceived the study, designed and interpreted the experiments, and wrote the manuscript with R.M.L. S.K., S.V., H.E.L., M.L., and C.E.O performed experiments and provided expertise. A.W.S. and C.E.O. helped with analysis of scRNA-seq data; N.K. helped with analysis of the ATAC-seq data. R.R.-G., O.D.K. and M.S.D. provided specialized reagents and contributed oversight and review. R.M.L. directed the study and wrote the paper with V.S.C. with input from the co-authors.

## INCLUSION AND DIVERSITY

One or more of the authors of this paper self-identifies as an underrepresented ethnic minority in science. One or more of the authors of this paper received support from a program designed to increase minority representation in science.

## FIGURE LEGENDS

**Supplementary Figure 1.**
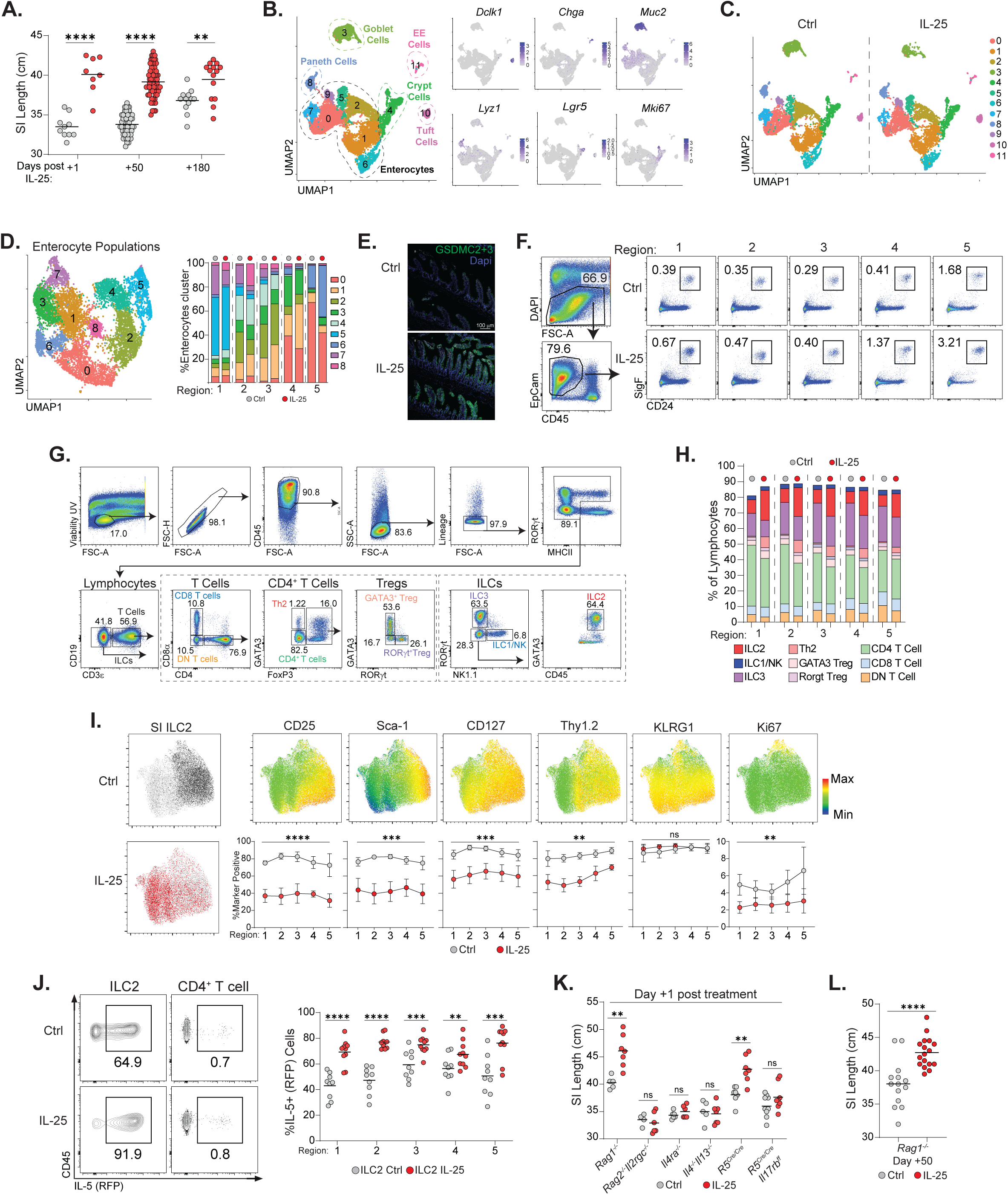
Small intestinal adaptation induced by IL-25 treatment, Related to Figure 1. (A) Small intestinal length at +1, +50, and +180 days after control or IL-25 treatment. (B) Epithelial cell UMAP generated by scRNA-seq and feature plots showing cell-type specific gene expression. (C) Epithelial cell UMAP from control or IL-25 treatment. (D) Enterocyte UMAP and abundance across small intestinal regions. (E) Gasdermin C 2+3 expression by intestinal epithelial cells. (F) Gating strategy and representative plots of tuft cells across small intestine. (G) Gating strategy and representative plots of lymphocytes populations in small intestine. (H) Abundance of lymphocytes populations in small intestine. (I) UMAP generated by flow cytometry of small intestinal ILC2s displaying phenotype based on treatment condition. Expression of indicated makers and quantification. (J) Representative plots of IL-5 (RFP) expression by ILC2s and CD4+ T cells in the small intestine; abundance of IL-5+ ILC2s. (K) Small intestinal length at day +1 after IL-25 treatment in mice from indicated genotypes. (L) Small intestinal length at +50 d after IL-25 treatment in *Rag1^-/-^* mice. Data pooled from at least two independent experiments (A, H, I, J, K, L) or from one experiment representative of at least 2 experiments (E, F). Each dot represents an individual mouse (A, J, K, L); Dot represents mean ± SD, n = 5-8 (J). Unpaired t test was performed. Statistical significance is indicated by *p < 0.05, **p < 0.01, ***p < 0.001, ****p < 0.0001; ns, not significant.

**Supplementary Figure 2.**
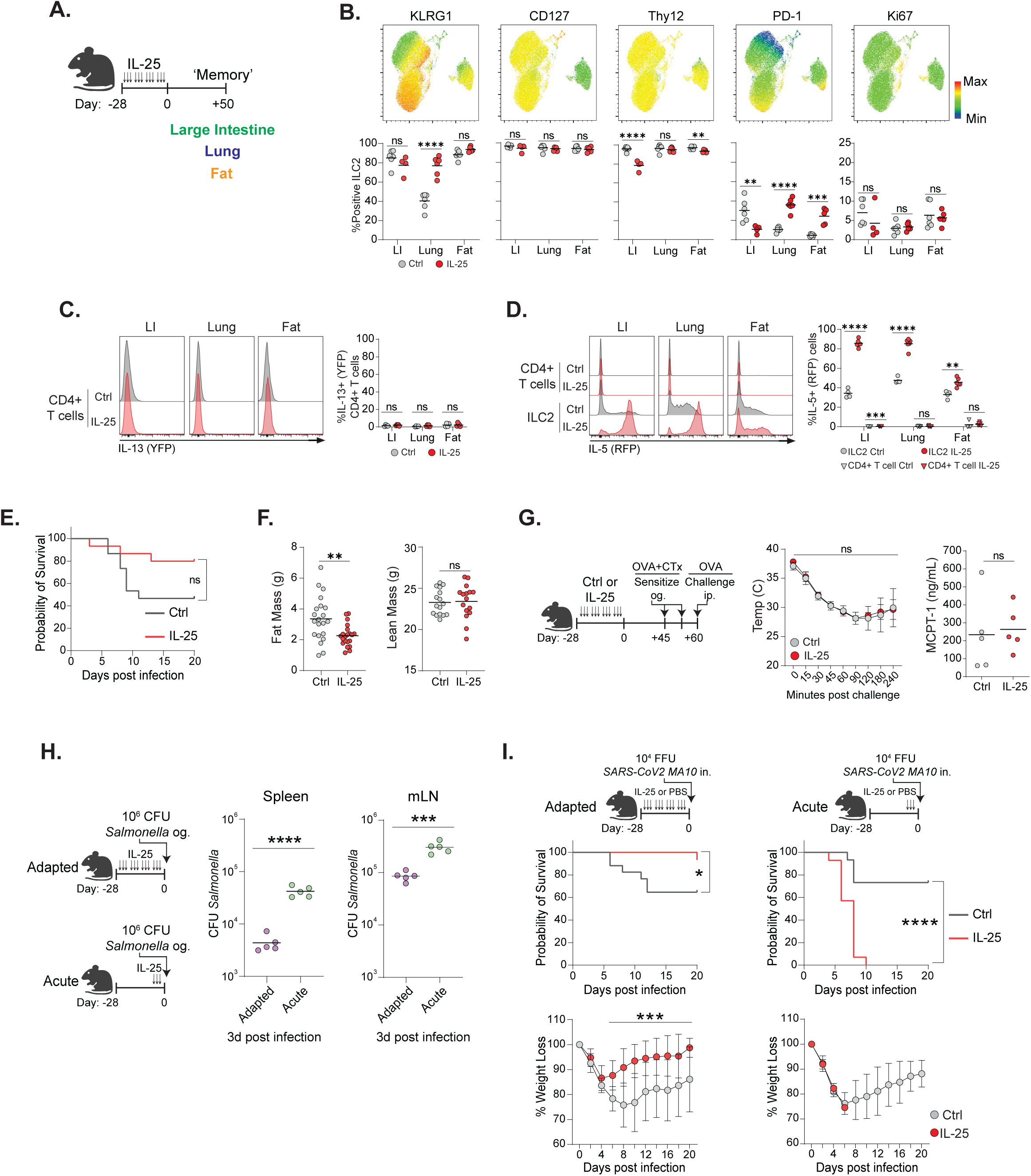
Alterations of ILC2 populations in tissues outside the small intestine and impact of IL-25 treatment on resilience to infectious challenges, adiposity, and food allergy, Related to Figure 2. (A) The IL-25 regimen and examination of the colon (LI), lung, and fat +50 d after treatment. (B) Expression and quantification of indicated markers by tissue ILC2s. (C) Representative histograms and quantification of IL-13 (YFP) expression by tissue CD4+ T cells. (D) Representative histograms and quantification of IL-5 (RFP) expression by tissue ILC2s and CD4+ T cells. (E) Survival during infection with SARS-CoV-2 MA10 in mice treated with the IL-25 regimen. (F) Fat and lean mass in mice at +1 d after control or IL-25 treatment. (G) Core body temperature of orally OVA-sensitized mice at indicated times after systemic OVA challenge. Serum MCPT-1 protein levels 60 minutes after challenge. (H) Titers of *Salmonella* in the spleen and mesenteric lymph nodes (mLN) +1 d after adapted or acute IL-25 treatment. (I) Survival and changes in weight during infection with SARS-CoV-2 MA10 in mice treated with the IL-25 adapted or acute regimen. Data pooled from at least two independent experiments (B, C, D, E, F, I) or from one experiment (G,H). Each dot represents an individual mouse (B, C, D, F, G, H); Dot represents mean ± SD, n = 5-15 (G, I). Unpaired t test (B, C, D, G, H), two-tailed t tests of area under the curve (weight-I), Long-rank (Mantel-Cox) test (survival-E,I), or two-way ANOVA (Bonferroni’s post-hoc test) (G) was performed. Statistical significance is indicated by *p < 0.05, **p < 0.01, ***p < 0.001, ****p < 0.0001; ns, not significant.

**Supplementary Figure 3.**
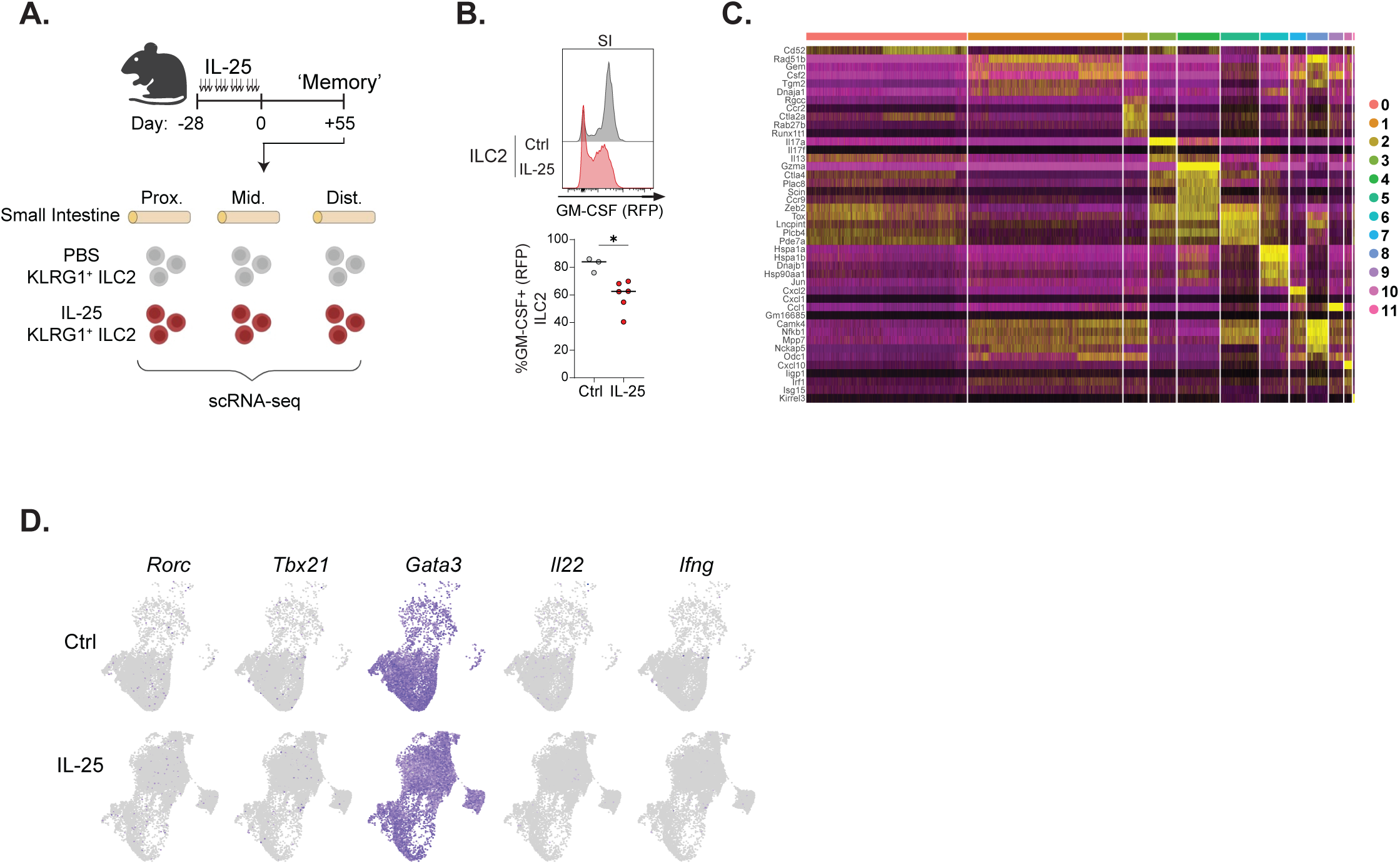
ILC2 transcriptional state after IL-25 conditioning, Related to Figure 3. (A) ILC2 isolation from the small intestines of control or IL-25 treated mice for scRNA-seq. (B) Representative histogram and quantification of GM-CSF (RFP) expression by small intestinal ILC2s. (C) Expression heatmap showing the top 5 markers for each ILC2 population. (D) Feature plots showing expressing on indicated genes on the ILC2 UMAP space. Data pooled from at least two independent experiments (B). Each dot represents an individual mouse (B). Unpaired t test was performed. Statistical significance is indicated by *p < 0.05.

**Supplementary Figure 4.**
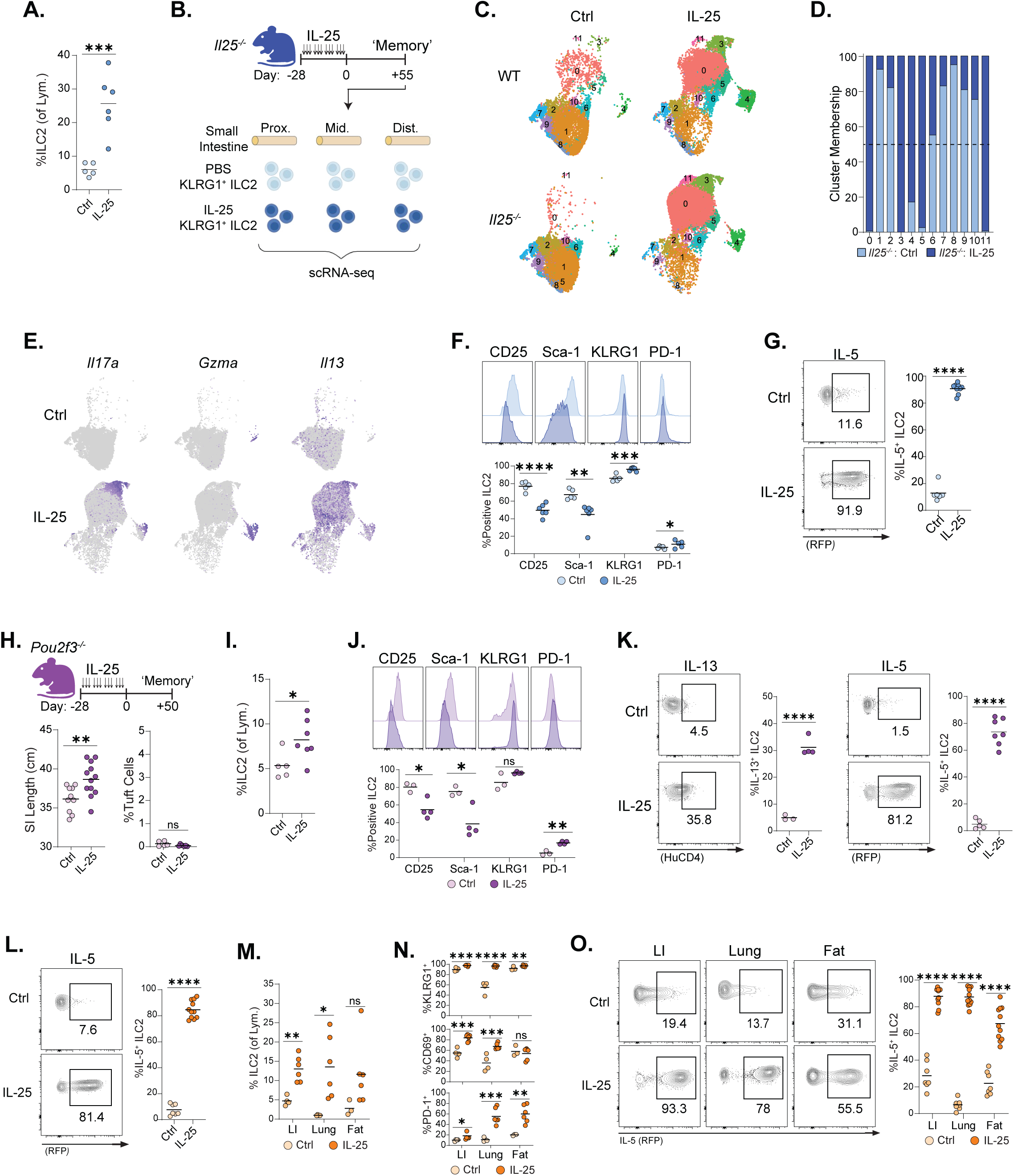
ILC2 memory features are maintained independently from alarmins and tuft cells, Related to Figure 4. (A) ILC2 abundance in the small intestine of *Il25^-/-^* mice after the IL-25 regimen. (B) ILC2 isolation from the small intestine of control or IL-25 treated *Il25^-/-^* mice for scRNA sequencing. (C) UMAP generated by scRNA-seq showing ILC2 populations from control or IL-25 treated WT or *Il25^-/-^* mice. (D) ILC2 cluster representation in *Il25^-/-^* mice. (E) Feature plots showing expression on indicated genes on the ILC2 UMAP space from *Il25^-/-^*mice. (F) Representative histograms and quantification of indicated markers by small intestine ILC2s from *Il25^-/-^* mice. (G) Representative plots and quantification of IL-5 (RFP) expression by small intestine ILC2s from *Il25^-/-^* reporter mice. (H) The IL-25 treatment regimen in *Pou2f3^-/-^* mice and quantification of small intestine length and tuft cell abundance. (I) ILC2 abundance in the small intestine of *Pou2f3^-/-^* mice. (J) Representative histograms and quantification of indicated markers by small intestine ILC2s from *Pou2f3^-/-^* mice. (K) Representative plots and quantification of IL-13 (HuCD4) and IL-5 (RFP) expression by small intestine ILC2s from *Pou2f3^-/-^* reporter mice. (L) Representative plots and quantification of IL-5 (RFP) expression by small intestine ILC2s from *Il25^-/-^Il1rl1^-/-^Crlf2^-/-^Il18^-/-^* reporter mice. (M) ILC2 abundance in tissues of *Il25^-/-^Il1rl1^-/-^Crlf2^-/-^Il18^-/-^* mice after IL-25 treatment. (N) Quantification of indicated markers by tissue ILC2s from *Il25^-/-^Il1rl1^-/-^Crlf2^-/-^Il18^-/-^* mice. (O) Representative plots and quantification of IL-5 (RFP) expression by tissue ILC2s from *Il25^-/-^ Il1rl1^-/-^Crlf2^-/-^Il18^-/-^* reporter mice. Data pooled from at least two independent experiments (A, F, G, H, I, J, K, L, M, N, O). Each dot represents an individual mouse (A, F, G, H, I, J, K, L, M, N, O); Unpaired t test was performed. Statistical significance is indicated by *p < 0.05, **p < 0.01, ***p < 0.001, ****p < 0.0001; ns, not significant.

**Supplementary Figure 5.**
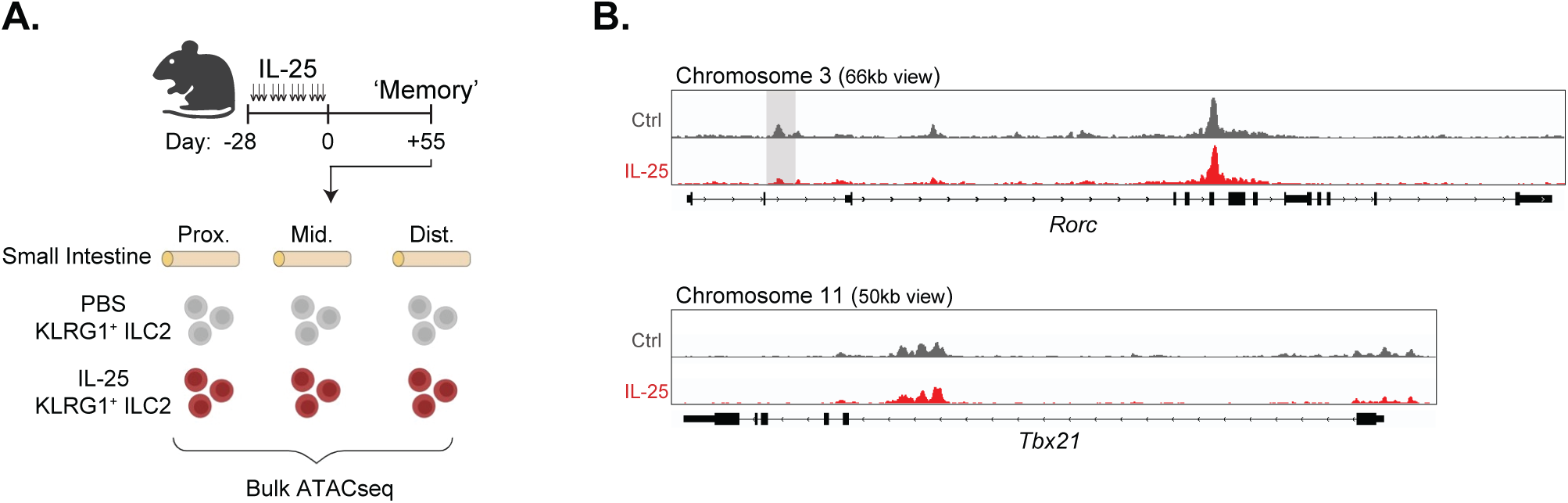
ATAC-seq of intestinal ILC2s, Related to Figure 5. (A) ILC2 isolation from the small intestines of control or IL-25 treated mice for ATAC-seq. (B) Genomic tracts of the *Rorc* gene and the *Tbx21* gene. Shaded areas represent regions with significantly less peaks (grey) in IL-25 conditioned ILC2s.

**Supplementary Figure 6.**
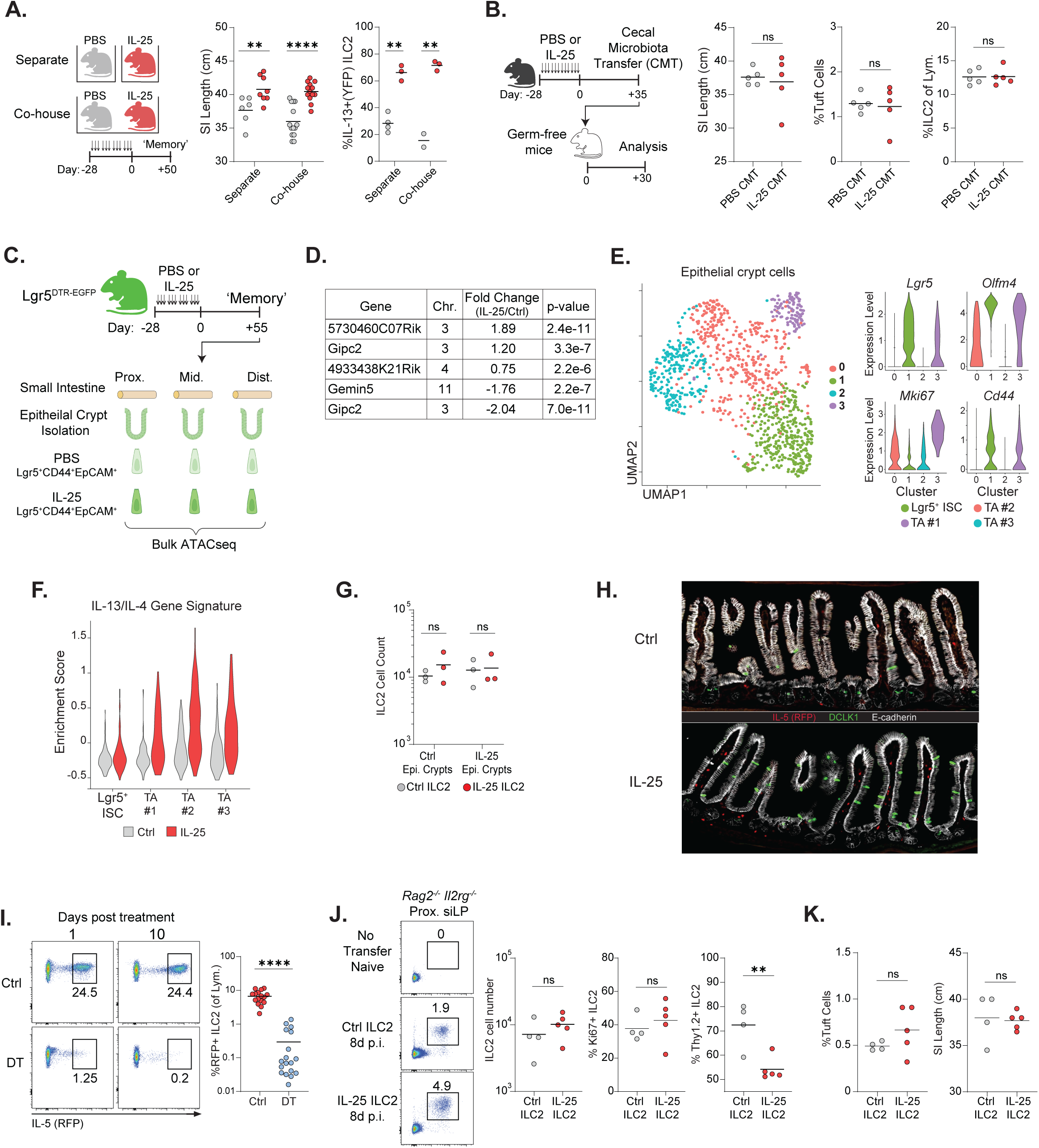
Dissecting adaptation in the small intestine, Related to Figure 6. (A) Small intestinal length and IL-13+ intestinal ILC2s in reporter mice. Mice were co-housed or separately housed since birth and maintained until analysis at +50 days. (B) Small intestinal length, tuft cell abundance, and intestinal ILC2 abundance in germ-free mice which received cecal microbiota transfer from control or IL-25 treated mice. (C) Lgr5+ ISC isolation from the small intestines of control or IL-25 treated mice for ATAC-seq. (D) Differentially expressed peaks between Lgr5+ intestinal stem cells from control or IL-25 treated mice. (E) Epithelial crypt UMAP generated by scRNA-seq and violin plots showing expression of genes used to identify epithelial crypt cell populations. (F) Violin plot showing expression of the IL-13 + IL-4 signature genes amongst epithelial crypt cell populations in control or IL-25 treated mice. (G) Absolute numbers of ILC2s after 6 days of co-culture with epithelial crypt organoids. (H) Representative image of IL-5 expressing cells and DCLK1+ tuft cells within the intestines of control or IL-25 treated mice. (I) Representative plots and quantification of ILC2 (RFP+) abundance in the small intestine of ILC2-DTR mice 1- or 10-days after treatment with PBS (Control) or diphtheria toxin (DT). (J) Representative plots and quantification of small intestinal ILC2 abundance, Ki67+ ILC2s, and Thy1.2+ ILC2s in *Rag2^-/-^Il2rg^-/-^* recipient mice 8 days after *N. brasiliensis* infection. (K) Tuft cell abundance and small intestinal length in *Rag2^-/-^Il2rg^-/-^* recipient mice 8 days after *N. brasiliensis* infection. Data pooled from at least two independent experiments (A, H, I, J, K), from one experiment (B), or from one representative experiment (G). Each dot represents an individual mouse (A, B, H, I, J, K); each dot is the average of 3 technical replicates (H). Unpaired t test was performed. Statistical significance is indicated by **p < 0.01, ****p < 0.0001; ns, not significant.

## RESOURCE AVAILABILITY

### Lead contact

Further information and requests for reagents may be directed to, and will be fulfilled by, the lead contact Richard Locksley.

### Materials availability

Reagents generated or used in this study are available on request from the lead contact with a completed Materials Transfer Agreement. Information on reagents used in this study is available in the key resources table.

## EXPERIMENTAL MODEL AND SUBJECTIVE DETAILS

### Mice

Wild-type C57BL/6J (000664) and *Rag1^-/-^* (B6.129S7-*Rag1^tm1Mom^*/J; 002216) mice were purchased from the Jackson Laboratory. *Rag2^-/-^Il2rg^-/-^* (C57BL/6NTac.Cg-*Rag2^tm1Fwa^ Il2rg^tm1Wjl^*; 4111-F) mice were purchased from Taconic Biosciences. B6.*Il13^YetCre13^*, B6.*Il5^Red5^*, B6.*Il13^Smart^*, B6.*Csf2^tdTomato^* reporter alleles on C57BL/6J (B6) backgrounds were bred and maintained as described (65, 22, 66, 67). B6.*Il4ra^-/-^*and B6.*Il4^-/-^Il13^-/-^* were obtained or generated and maintained as previously described (11, 66). B6.*Il25^-/-^* (68) were bred to *Il5^Red5^*and *Il13^Smart^* reporter alleles. B6.*Pou2f3^-/-^*mice were provided by M. Anderson and were bred to *Il5^Red5^* and *Il13^Smart^* reporter alleles. Il25^-/-^, Il1rl1^-/-^ (69), Crlf2^-/-^ (70) and Il18^-/-^ mice (B6.129P2-*Il18^tm1Aki^*/J; 004130) were intercrossed to generate *Il25^-/-^*, *Il1rl1^-/-^*, *Crlf2^-/-^*, *Il18^-/-^* quadruple-deficient mice expressing *Il5^Red5^* and *Il13^Smart^* reporter alleles as described (71). *Il17rbf/f* mice were provided by U. Siebenlist (72) and crossed to *Il5^Red5^*mice. ROSA26-iDTR (C57BL/6-Gt(ROSA)26Sor*^tm1(HBEGF)Awai^*/J; 007900) mice were obtained from the Jackson Laboratory and crossed to *Il5^Red5^*mice. R26-CreERT2 (B6.129-Gt(ROSA)26Sor*^tm1(cre/ERT2)Tyj^*/J; 008463) were obtained from the Jackson Laboratory and crossed to the *Il4ra^ff^* mice provided by A. Chawla (73). *Lgr5^DTR-EGFP^* mice were provided by O. Klein (74). For all experiments, sex-matched mice aged 7-16 weeks were used to begin the regimens. Mice were maintained under specific pathogen-free conditions. All animal procedures were approved by the UCSF Institutional Animal Care and Use Committee.

### IL-25 treatment

Mice were injected with 500 ng IL-25 (Peprotech or R&D) intraperitoneally 3 times a week for four weeks, for 12 total injections. Control mice were injected with an equal volume of PBS. All injections were given in 200 μl. The ceca of all mice were checked for the presence of *Tritrichomonas*. In general, C57BL/6J mice were *Tritrichomonas* free and all genetically modified strains were *Tritrichomonas* colonized. For some experiments, all mice in a cage were injected with either IL-25 or PBS. For other experiments, mice in the same cage were given either IL-25 or PBS and kept co-housed until analysis.

### Intestine length

Promptly after euthanasia, the proximal small intestine was cut at the pyloric sphincter, carefully pulled out to avoid tearing, and then cut distally where it meets the cecum. The mesenteric fat was completely removed, and the intestine held in the air from the distal portion for 5 seconds to allow the tissue to reach its full length. The tissue was then held by the proximal region in the air and measured vertically alongside a measuring tape.

### Infections, food allergy, and cecal microbiota transfer

For *Nippostrongylus brasiliensis* (*N. brasiliensis*) infection, mice were infected subcutaneously with 500 (*Il25^-/-^ Il1rl1^-/-^ Crlf2^-/-^ Il18^-/-^* mice) or 250 (*Rag1^-/-^*or *Rag2^-/-^Il2rg^-/-^* mice) L3 worms or orally gavaged with 150 (*Rag1^-/-^* mice) L5 worms. Mice were euthanized at indicated times and intestinal worm burden quantified as described (22). For *Salmonella* Typhimurium SL1344 infection, C57BL/6J mice were fasted for 4 hours, then pretreated with streptomycin by oral gavage, and 24 hours later inoculated with 10^6^ CFU of bacteria by oral gavage. Mice were sacrificed on day 3 post infection and bacterial burdens quantified in the spleens and mesenteric lymph nodes. For SARS-CoV-2 (MA-10) infection (75), 24-week-old male C57BL/6J mice were challenged with 10^4^ FFU of MA-10 by intranasal administration in 50 μL of PBS. Mice were weighed before and every other day after infection until day 20 post-infection. Survival was monitored daily. SARS-CoV-2 strain (MA-10) was propagated and tittered in Vero-hACE2-TMPRSS2 cells as previously described (76) and all virus infection experiments were performed in approved biosafety level 3 facilities with appropriate positive-pressure respirators, personal protective gear, and containment at Washington University in St. Louis. For food allergy (28), C57BL/6J mice were sensitized by oral gavage with 200 μl of 250 mg/mL albumin + 10 μg cholera toxin in 5% NaHCO3^-^ solution on day 45 and 53 post treatment. Anaphylactic challenge was induced on day 60 by intraperitoneal injection of 50 μg EndoFit OVA. Mouse core body temperature was measured by rectal probe before challenge and at indicated time points after challenge. Blood was collected one hour post challenge and serum MCPT-1 levels determined by ELISA. For cecal microbiota transfer, cecal contents were isolated from two control and two IL-25 treated C57BL/6J mice at +50 days in an aerobic chamber. Cecal contents were resuspended in PBS and passed through a 100 μm filter to remove large debris. Germ-free mice were reconstituted with the cecal content suspension and examined 35 days later.

### Weight and body mass assays

C57BL/6J male mice, aged 7-8 weeks, were weighed prior to IL-25 treatment, once a week during treatment, and every two weeks following treatments. Fat and lean mass were measured 2-3 days after the final injection of PBS or IL-25 by MRI using an EchoMRI (Echo Medical Systems LTD) following the manufacturer’s instructions.

### Tissue preparation

Small intestinal single cell suspensions were prepared either from all 5 regions (where indicated) or the most distal region #5, as previously described (77). Briefly, small intestine length was measured, and the tissue cut into 5 equal-length pieces. Luminal contents were flushed out with PBS, Peyer’s patches removed, and tissue cut open. Pieces were incubated with epithelial cell extraction buffer (EEB: Ca2+/Mg2+ free HBBS, 3% FCS, 10 mM HEPES, 10 mM EDTA, 10 mM DTT) at 37℃ with gentle shaking for 15 minutes. Tubes were then vortexed at full speed for 30 seconds and supernatants poured through a 100 μm filter and collected for epithelial cell analysis. The tissue was placed with fresh EEB at 37℃ for 15 minutes with gentle shaking, then vortexed for 30 seconds and supernatant discarded. Tissue was placed in rinse buffer (Ca2+/Mg2+ sufficient HBBS, 3% FCS, 10 mM HEPES) and incubated at 37℃ with gentle shaking for 5 minutes. Tissue was then placed in GentleMACS C tubes (Miltenyi Biotec) with digestion buffer (DB: Ca2+/Mg2+ sufficient HBBS, 3% FCS, 10 mM HEPES, 30 μg/mlDNase I, 0.1 Wunsch/ml Liberase TM), minced with scissors, and incubated at 37℃ with gentle shaking for 10-15 minutes. Single-cell suspensions were obtained using a gentleMACS tissue dissociator (Miltenyi Biotec), running program m_intestine_0, and then pouring through a 100 μm filter and washing with FACS buffer (PBS, 3% FCS). Large intestine single cell suspensions were prepared similarly to the small intestine, except that the whole tissue length was used. For lung and fat single cell suspensions, the left lung lobe or perigonadal visceral adipose tissue was isolated and placed in a GentleMACS C tube with digestion buffer, minced with scissors, incubated at 37℃ with gentle shaking for 20 (lung) or 40 (fat) minutes, the gentleMACS tissue dissociator program lung_02 was used, and the suspension then poured through a 100 μm filter and washed with FACS buffer.

### Flow cytometry and cell sorting

Single-cell suspensions were incubated with Fc Block and staining antibodies to surface markers; for spectral cytometry experiments Brilliant Stain Buffer was used. Cells were kept at 4℃ throughout the staining procedure, and DAPI was used to identify live cells which were enumerated by flow cytometry with CountBright Absolute Counting Beads. For intracellular staining of transcription factors, the FoxP3/Transcription Factor Staining Buffer Set was used and fixable violet dead or fixable blue dead stain used to exclude dead cells. Samples were analyzed by FACS on a 5-laser LSRFortessa X-20 (BD Bioscience) or a 5-laser Cytek Aurora (Cytek). Data were analyzed with FlowJo v10.8.0. ILC2 were sorted from the proximal, mid, and distal small intestine, and then combined for a given sample. ILC2 were sorted as KLRG1+Lineage-(CD3, CD4, CD8a, CD19, NK1.1, NKp46, CD11b, CD11c, F480, Gr-1, Ter-119) CD45+ cells. ILC2 were sorted to >90% purity using a MoFlo XDP (Beckman Coulter).

### Immunofluorescence (IF) and whole mount imaging

For IF, the small intestine lumen was flushed with PBS, cut open, and fixed in 4% paraformaldehyde for 4 hours at room temperature or overnight at 4℃. Tissue was washed with PBS and incubated with 30% (w/v) sucrose in PBS overnight at 4℃. Tissue was Swiss rolled and embedded in a mixture of 1:1 Tissue-Tek O.C.T and 30% sucrose and stored at –80°C until further processing. Tissue sections (8-10 μm) were cut on a Cryostat (Leica). Cryosections were dried for 15 minutes, washed 3x in PBS, incubated for 10 minutes in PBST (PBS + 0.1% Triton X-100), and blocked for 1 hour at room temperature in Antibody Dilution Solution (ADS: 1x Animal Free Block, 0.1% SDS, 0.1% Triton-X100 in H2O) containing 2.5% Donkey or Goat serum. Sections were incubated in ADS with primary antibodies for 2-24 hours, washed 3x with PBST, incubated in ADS with secondary antibody for 1-2 hours at room temperature, washed 3x in PBST, incubated 5 min with a 1:1000 dilution of DAPI (Stock 5 mg/ml in PBS), washed 3x in PBST, and mounted in Mounting Media or Prolong Gold and images acquired using a Zeiss immunofluorescence AxioImager M2 upright microscope with a Plan Apochromat 20X/0.8NA air objective running AxioVision software (Zeiss) or an inverted Nikon A1R scanning confocal microscope with a Plan Apo λ 20X/0.75NA air objective running NIS Elements software (Nikon). For whole mounts, a 2-3 cm piece of the proximal and distal intestine was incubated with Cubic L at 37°C with shaking overnight, washed 3x with PBS, blocked for 2 hours in 5% Donkey Serum in Whole Mount Blocking Buffer (WMBB: 1x Animal Free block, 1% NaN3, 0.3% Triton X-100 in PBS), incubated with primary antibodies in WMBB overnight at 4℃. Samples were washed 3x with PBS, incubated in secondary antibody and DAPI (1:500) in WMBB, washed 3x with PBS and incubated with Cubic-R for 2 hours at 37°C, followed by mounting in Cubic-R and imaging on a Zeiss LSM900 confocal microscope (10x water objective).

### Single cell RNA-seq

For the small intestine epithelial scRNA-seq, cells were isolated from C57BL/6J (a total of 2 IL-25 treated and2 PBS treated) mice 49 and 50 days after the last treatment. Cells were collected from both epithelial extraction buffer fractions and the epithelial cells (EpCAM+CD45-) were sorted to >95% purity. Sorted cells from a given region of PBS or IL-25 treated mice were labeled with TotalSeq anti-mouse Hashtag antibodies for 30 minutes at 4℃, followed by 3 washes with FACS buffer, and counting of cells. Equal number of cells from each region were combined and single-cell libraries from a total of 50,000 cells were prepared with the Chromium Next GEM Single Cell 3ʹ v3.1:Dual Index Libraries following the manufacturer’s protocol. For ILC2 scRNA-seq, cells were isolated from C57BL/6J (a total of 2 IL-25 treated and 2 PBS treated) mice and *Il25^-/-^* (a total of 2 IL-25 treated and 2 PBS treated) mice 55 days after the last treatment. ILC2 were sorted to >95% purity from the proximal, mid, and distal small intestine, and then combined for a give genotype and treatment condition. Single-cell libraries from a total of 50,000 cells per genotype and treatment condition were prepared with the Chromium Next GEM Single Cell 3’ GEM, Library and Gel Bead Kit v3.1. Libraries were sequenced on the NovaSeq 6000 at the UCSF Institute for Human Genetics. For scRNA-seq analysis of the small intestine epithelium, single-cell transcripts were aligned to the mouse genome and processed via the 10X Cell Ranger software version 4.0. The R package Seurat (version 4.1) was used for single-cell transcriptome analysis. The MULTIseqDemux function was used to demultiplex samples using the HTO CITE-seq assay data. Cells were removed according to the following thresholds: <200 genes/cell or >5000 genes per cell, >20000 UMIs/cell, 20% mitochondrial content, <0.01% hemoglobin read content. For the ILC2 scRNA-seq analysis, single-cell transcripts were aligned to the mouse genome and processed via the 10X Cell Ranger software version 7.0. The R package Seurat (version 4.3) was used for single cell transcriptome analysis. Cells were removed according to the following thresholds: <1000 genes/cell or >4000 genes per cell, >30000 UMIs/cell, 10% mitochondrial content, <0.001% hemoglobin read content to remove red blood cells. Among the epithelial or ILC2 cells retained, the effects of mitochondrial and ribosomal content were regressed out using the SCTransform normalization method prior to clustering. Genes were excluded from the final dataset if they were expressed in fewer than 3 cells. For differential expression analysis between clusters, genes were detected if they are expressed in at least 10% of cells in a cluster with a log fold change of at least 0.25. For DEG analysis between comparison groups, genes were detected if expressed in at least 5% of cells in the group with a log fold change of at least 0.25.

### ATAC-seq

For ILC2 ATAC-seq, cells were isolated from C57BL/6J (a total of 3 IL-25 treated and 3 PBS treated) mice 50-55 days after the last treatment. ILC2 were sorted to >95% purity from the proximal, mid, and distal small intestine, and combined for a given genotype and treatment condition. For epithelial stem cell ATAC-seq, GFP+CD44+CD45-cells were sort purified from intestinal crypts isolated from proximal, mid, and distal small intestines of *Lgr5^DTR-EGFP^* (a total of 3 IL-25 treated and 3 PBS treated) mice 50-60 days after the last treatment. Cells were subjected to Assay for Transposase-Accessible Chromatin using Sequencing (ATAC-Seq) (78). 30,000–55,000 cells were sorted into PBS + 10% FCS, washed, and lysed in 100 μl detergent (100 mM Tris-HCl pH 7.4, 10 mM NaCl, 3 mM MgCl_2_, 0.1% IGEPAL CA-630) on ice for 5 min to release the nuclei. Pelleted nuclei were incubated for 30 min with Tagment DNA Enzyme transposase. The reaction was stopped and DNA purified using the MinElute Reaction Cleanup kit. Eluted DNA was amplified with the NEBNext High-Fidelity 2x PCR Master Mix using Nextera index primers. The indexed libraries were purified using the MinElute PCR Reaction Cleanup before sequencing on an Illumina HiSeq 2500 with 25 bp paired-end reads. Raw reads were mapped to the mouse mm10 genome assembly using STAR alignment (--outFilterMismatchNoverLmax 0.04 --outFilterMismatchNmax 999 --alignSJDBoverhangMin 1 --outFilterMultimapNmax 1 --alignIntronMin 20 --alignIntronMax 1000000 --alignMatesGapMax 1000000). Bam files were generated by STAR. PCR duplicates removed by Picard, and peak calling performed using MACS2 (--keep-dup 1 --bw 500 -n output --nomodel --extsize 400 --slocal 5000 --llocal 100000 -q 0.01). Differential binding peaks were identified with DiffBind (method=DBA_DESEQ2), and peaks annotated using ChIPSeeker. To generate bigwig files for ATACseq datasets, all aligned bam files were merged by condition using samtools merge. Samtools view was then run to convert alignments to counts and bedtools genomecov was run to covert the merged bam files into a bedgraph files. Finally, bedGraphToBigWig (ucsc-tools/363) was used to generate the bigwig files displayed on browser tracks using the IGV browser.

### Organoid culture

Intestinal organoids were generated as described (79). Briefly, the intestine length was measured, the tissue divided into three equidistant segments, and the middle 5 cm of each segment taken for crypt isolation. Tissue was cut open and rinsed twice in PBS + Pen/Strep on ice, then incubated in Harvest Buffer (PBS + Pen/Strep + 2 mM DTT + 10 μM Y-27632+ 1 mM EDTA) with rocking on ice for 15 minutes. After 15 minutes, the tissue was shaken for 1 minute, transferred to Crypt Dissociation Buffer (PBS + Pen/Strep + 2 mM DTT + 10 μM Y-27632+ 5 mM EDTA), and incubated for 1 hour with gentle rocking on ice. The tissue was manually shaken for 1-3 minutes, filtered through a 70 µm filter, rinsed with 5 mL of ice cold Basal Medium (Advanced DMEM/F12 High Glucose, N-acetyl-L-cysteine, Pen/Strep, Glutamax, HEPES) and enumerated. 1000 crypts were plated per well in preheated 24-well plates in 50 μL Matrigel domes in ENR medium (Basal Medium, B27 Supplement, N2 Supplement, hrEGF (50 ng/mL), Noggin (100 ng/mL), and R-spondin conditioned medium). For experiments with IL-13 treatment of organoids, ENR medium + IL-13 (20 ng/mL) was added 3 days after plating. The medium was exchanged for fresh ENR + IL-13 at day 5. 6 days after plating, matrigel domes were washed with 1x PBS, then incubated with 0.5 pre-warmed TrypLE for ∼10 minutes, with trituration to promote organoid dissociation. The reaction was stopped with 1 mL of Basal medium + 10% FBS and the resulting cell suspensions washed and tuft cell abundance assessed by flow cytometry. For ILC2-organoid co-culture experiments, organoids were established from a 2 cm segment of the proximal duodenum of C57BL/6J mice. Intestinal ILC2 were sorted purified and 10,000 ILC2 /0.5 mL in ENR medium with 2-mercaptoethanol, IL-2 (10 ng/mL), and IL-7 (10 ng/mL) were added to wells with epithelial crypts set in matrigel. The medium was changed on day 3; ILC2s were spun down and resuspended in fresh medium and added back to their original well. For IL-13/IL-4 blocking studies, organoids were generated from *Lgr5^DTR-EGFP^* mice and plated with ILC2 and control IgG (10 μg/mL), or anti-IL-13 (100 ng/mL) and anti-IL-4 (10 μg/mL) antibodies, as indicated. Organoids were dissociated to single cell suspensions and analyzed by flow cytometry. Each experiment was repeated with n=3 technical replicates (organoid wells/condition) and 3-4 biological replicates (independent animals and experiments).

### ILC2 depletion

For ILC2 depletion experiments, *Red5^Cre/Cre^ x Rosa26^DTR/DTR^* mice were rested for at least 40 days after the final IL-25 treatment. They were injected with 250 ng diphtheria toxin intraperitoneally every other day for a total of 3 injections. Mice were analyzed 9-10 days after the final injection. For controls, *Red5^Cre/Cre^ x Rosa26^DTR/DTR^* mice were injected with PBS instead of diphtheria toxin, or IL-25 treated and rested *Red5^Cre/Cre^* mice were injected with diphtheria toxin.

### Tamoxifen induced deletion

For IL-4ra deletion, *Rosa26^CreERT2/CreERT2^ x Il4ra^ff^* were rested for at least 35 days after the final IL-25 treatment. They were put onto tamoxifen chow diets, and allowed to feed ad libitum for 3 weeks, and analyzed. For controls, *Rosa26^CreERT2/CreERT2^ x Il4ra^ff^* mice fed normal chow diets or IL-25 treated and rested *Il4ra^ff^* mice were placed on tamoxifen diets.

### ILC2 transfer to *Rag2 Il2rg*-deficent mice

ILC2s were isolated from C57BL/6J mice +30 days after the last treatment. ILC2s were sorted to >95% purity from the proximal, mid, and distal small intestine and combined for a given treatment condition. 1×10^5^ ILC2s were injected intravenously to each mouse, and mice were given 3 injections within a week, for a total of 3×10^5^ ILC2s per mouse. Recipient mice were rested for +30 days and infected with 250 L3 *N. brasiliensis*. Mice were kept on antibiotic water (100 μg polymyxin B/mL and 2 mg neomycin sulfate/mL) during infection and examined 8 days post infection.

### Quantification and statistical analysis

Experiments were performed using randomly assigned mice without investigator blinding. Data points represent individual animals pooled from multiple experiments. No data were excluded. Statistical significance was calculated as noted in the figures. No statistical methods were used to predetermine sample size. Statistical analyses were performed using Prism version 9.3.1 software (GraphPad Software, Inc.).

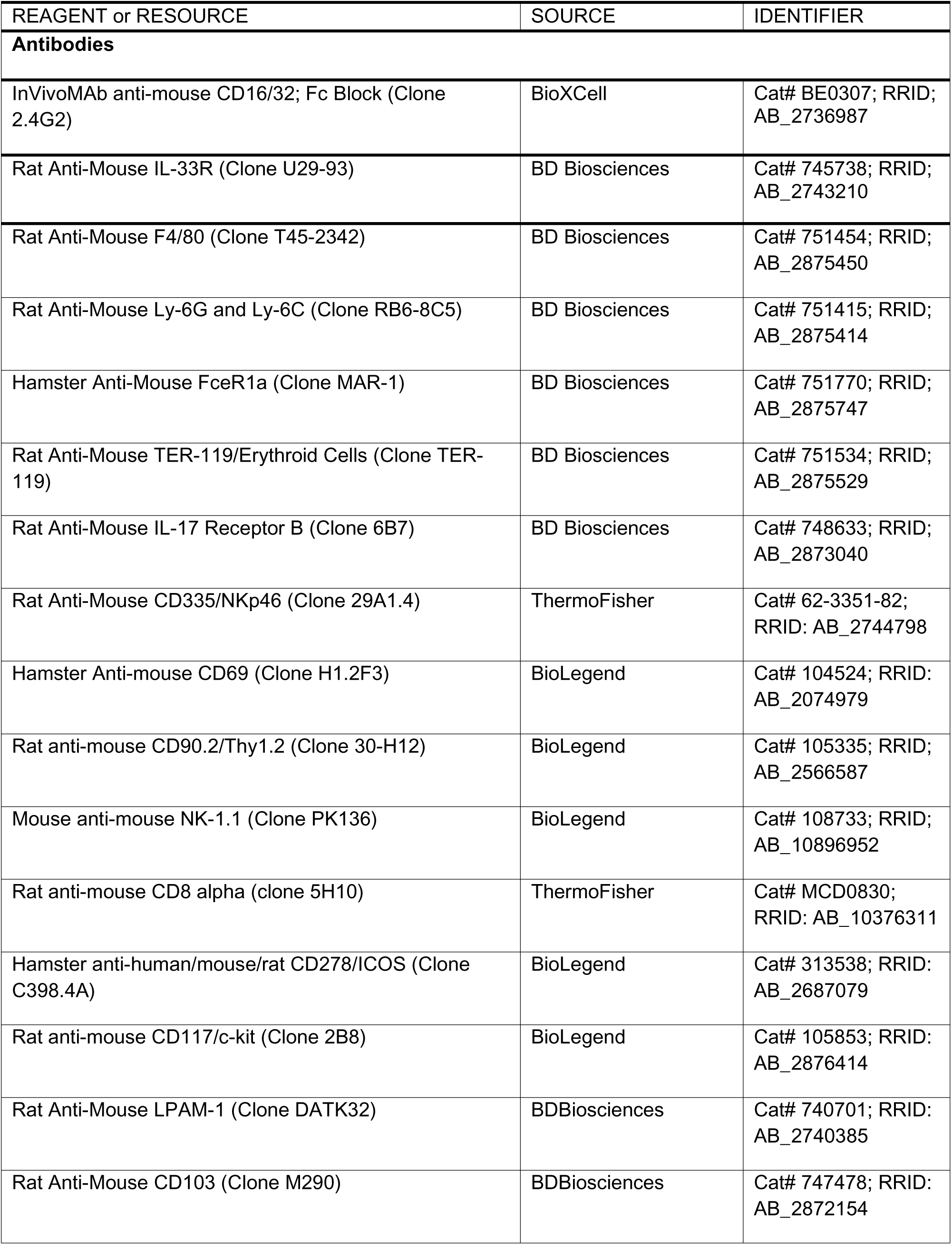
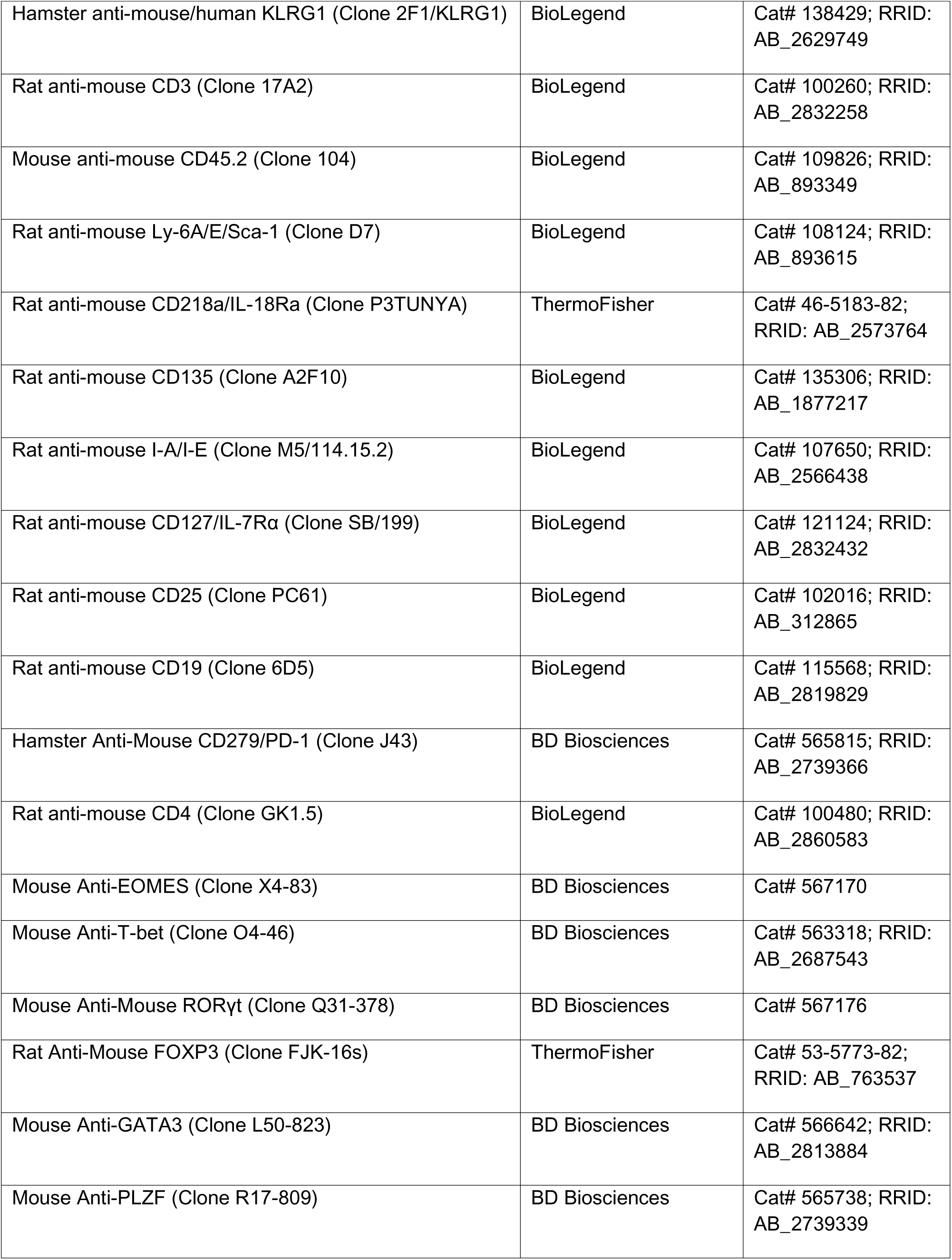
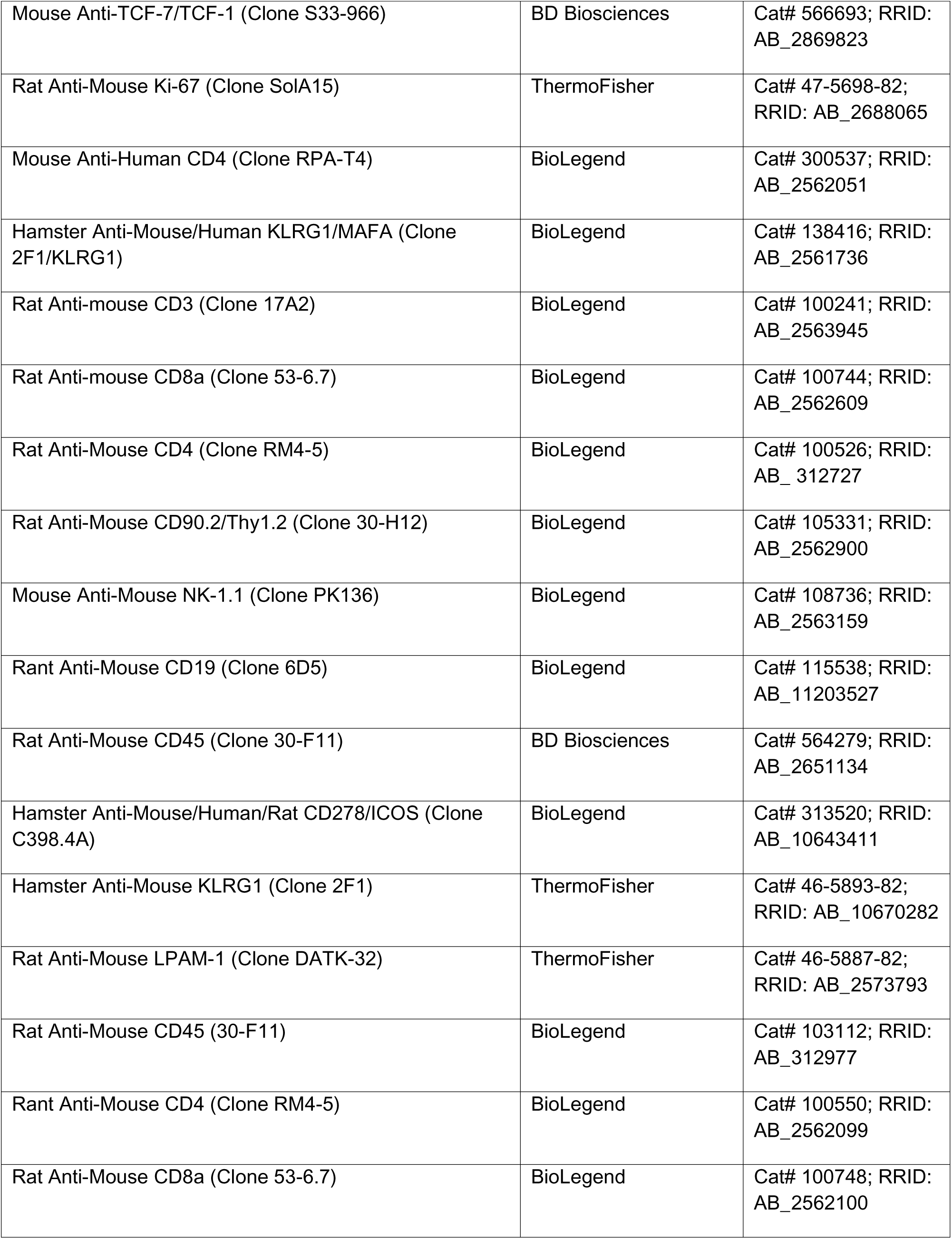
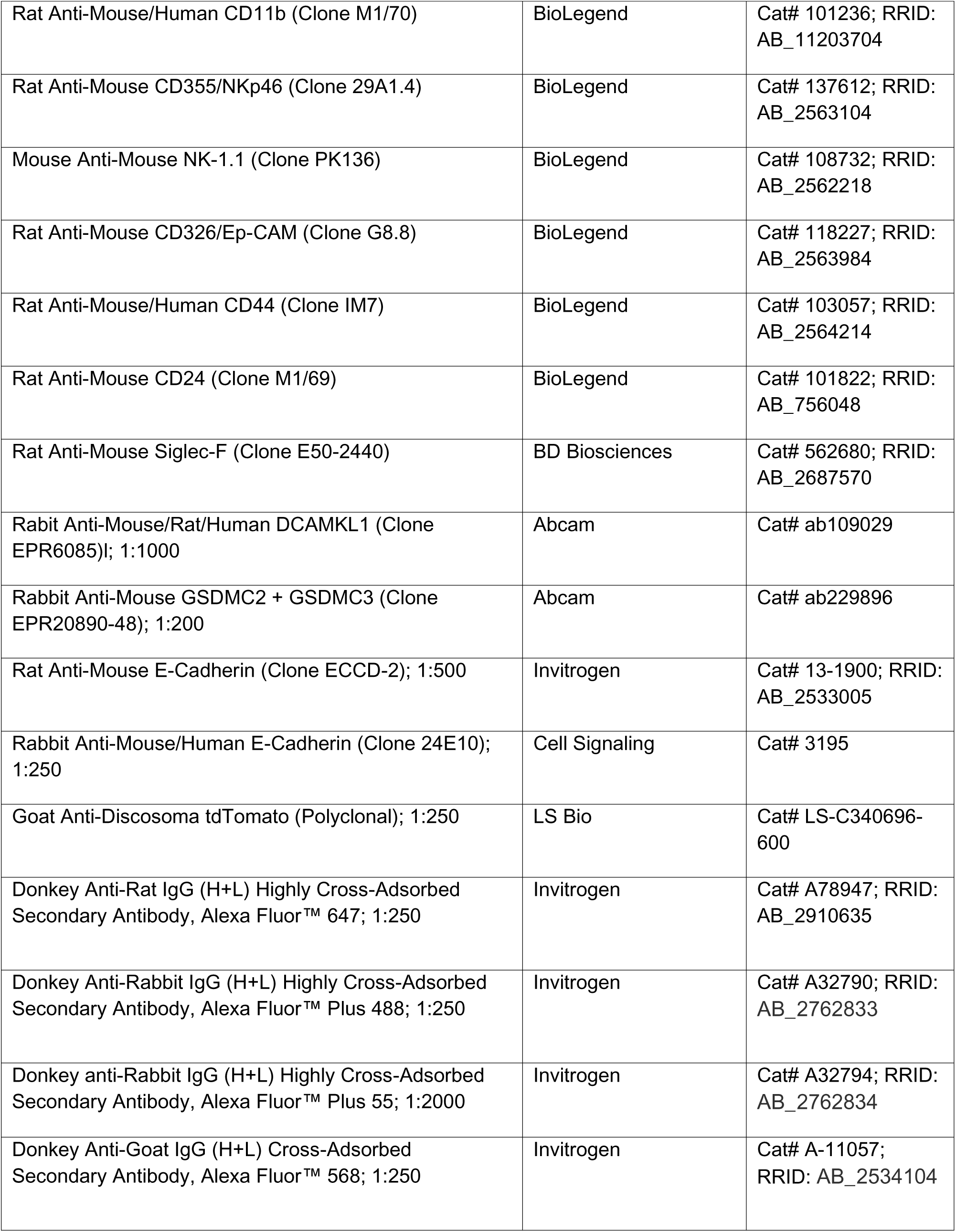
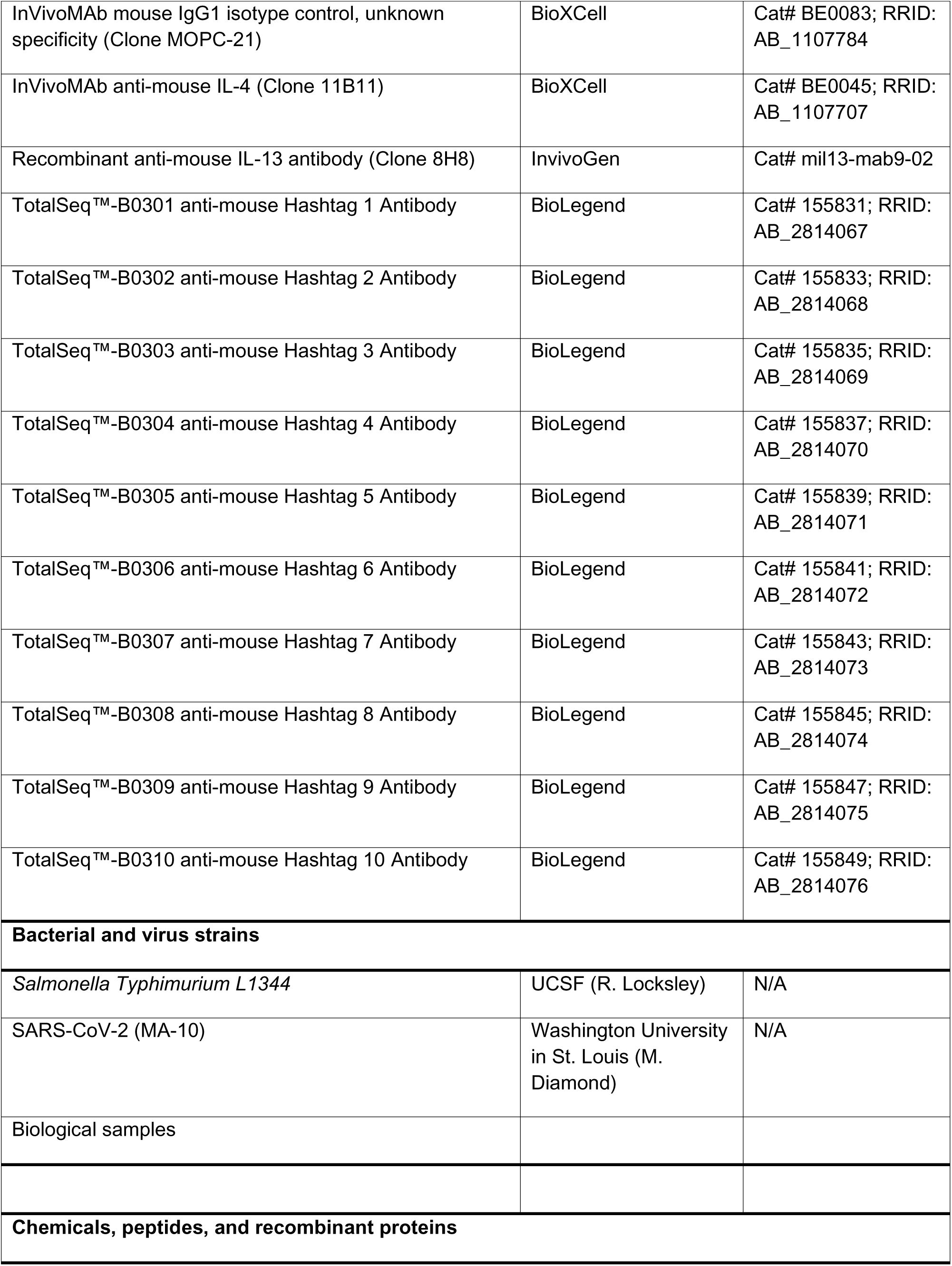
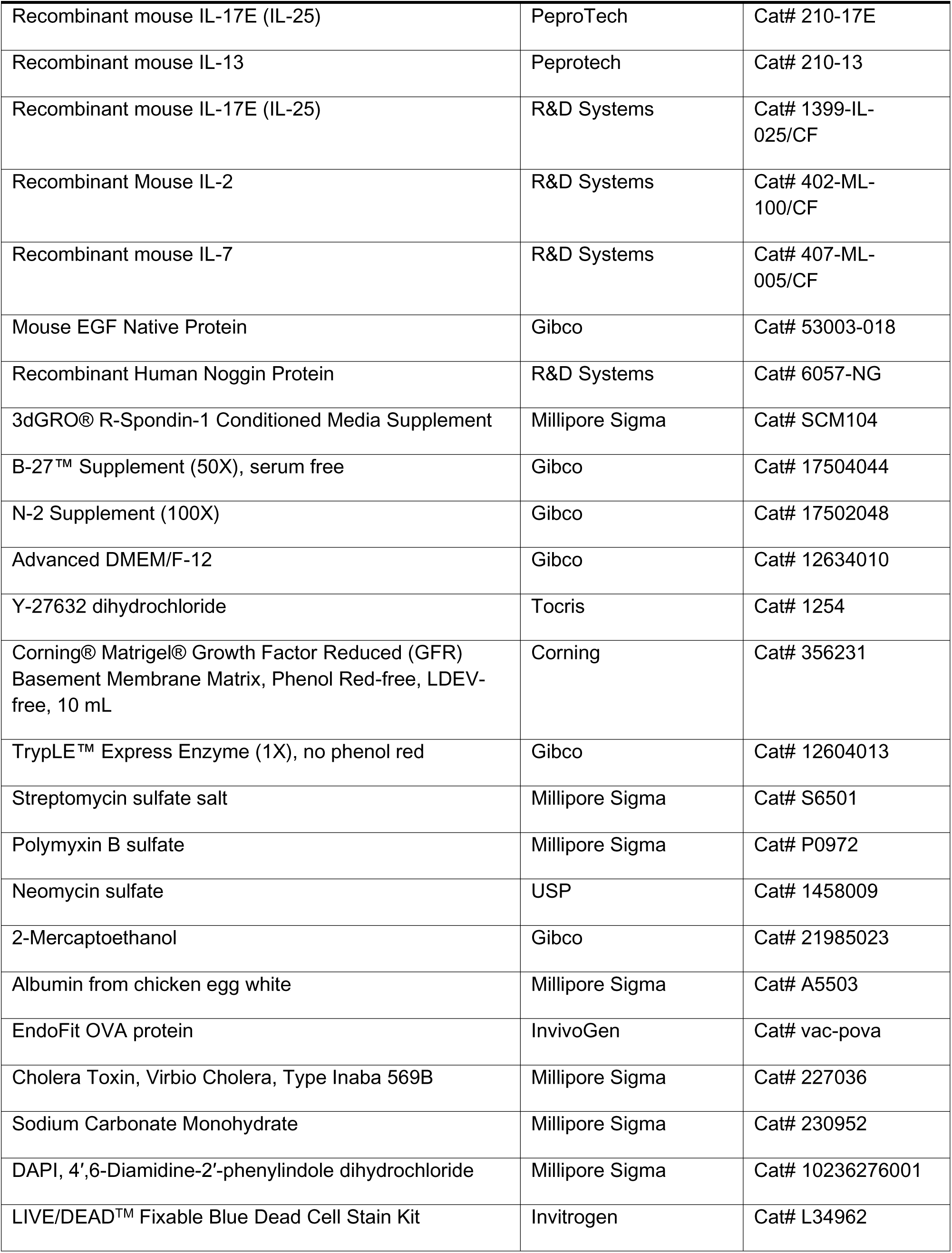
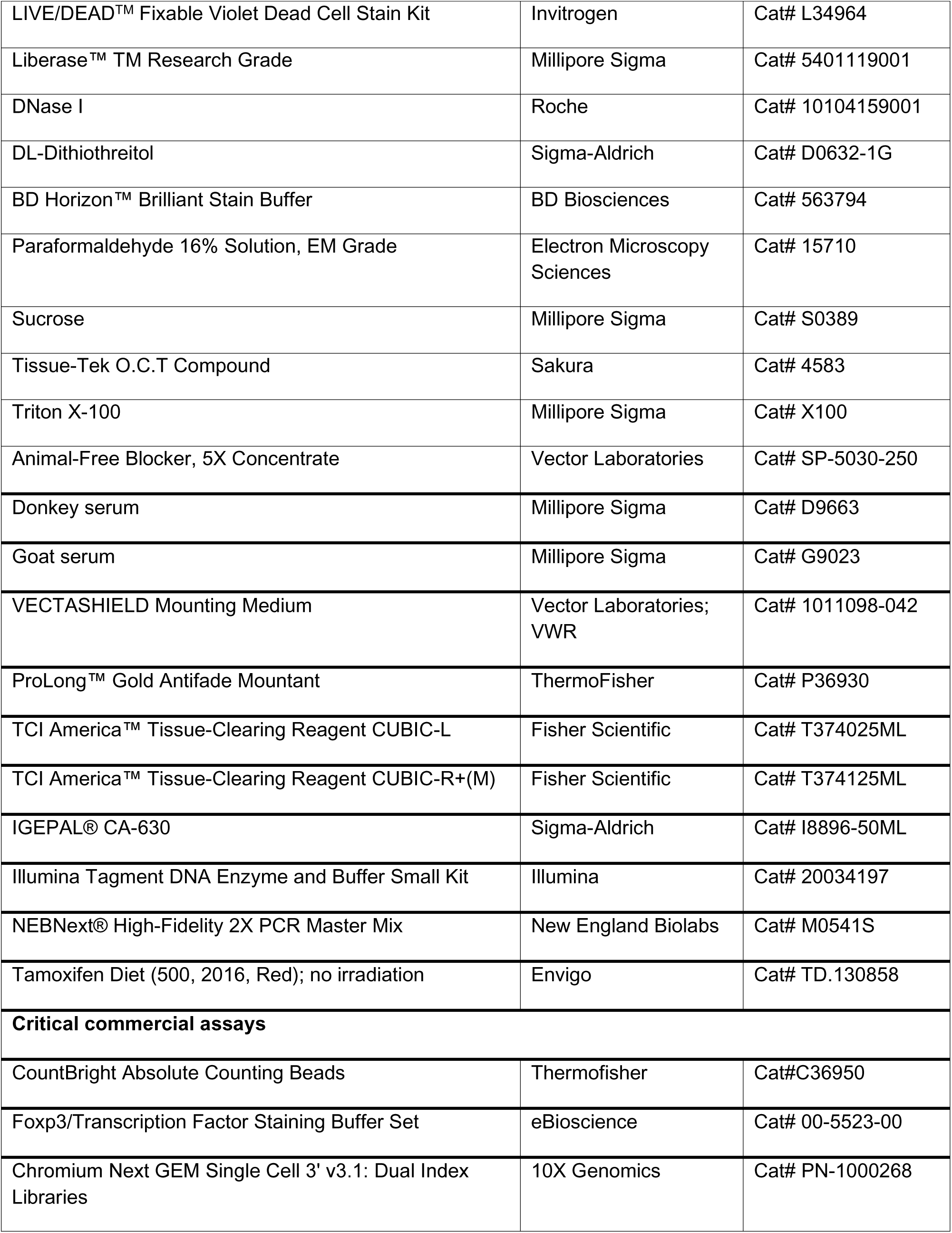
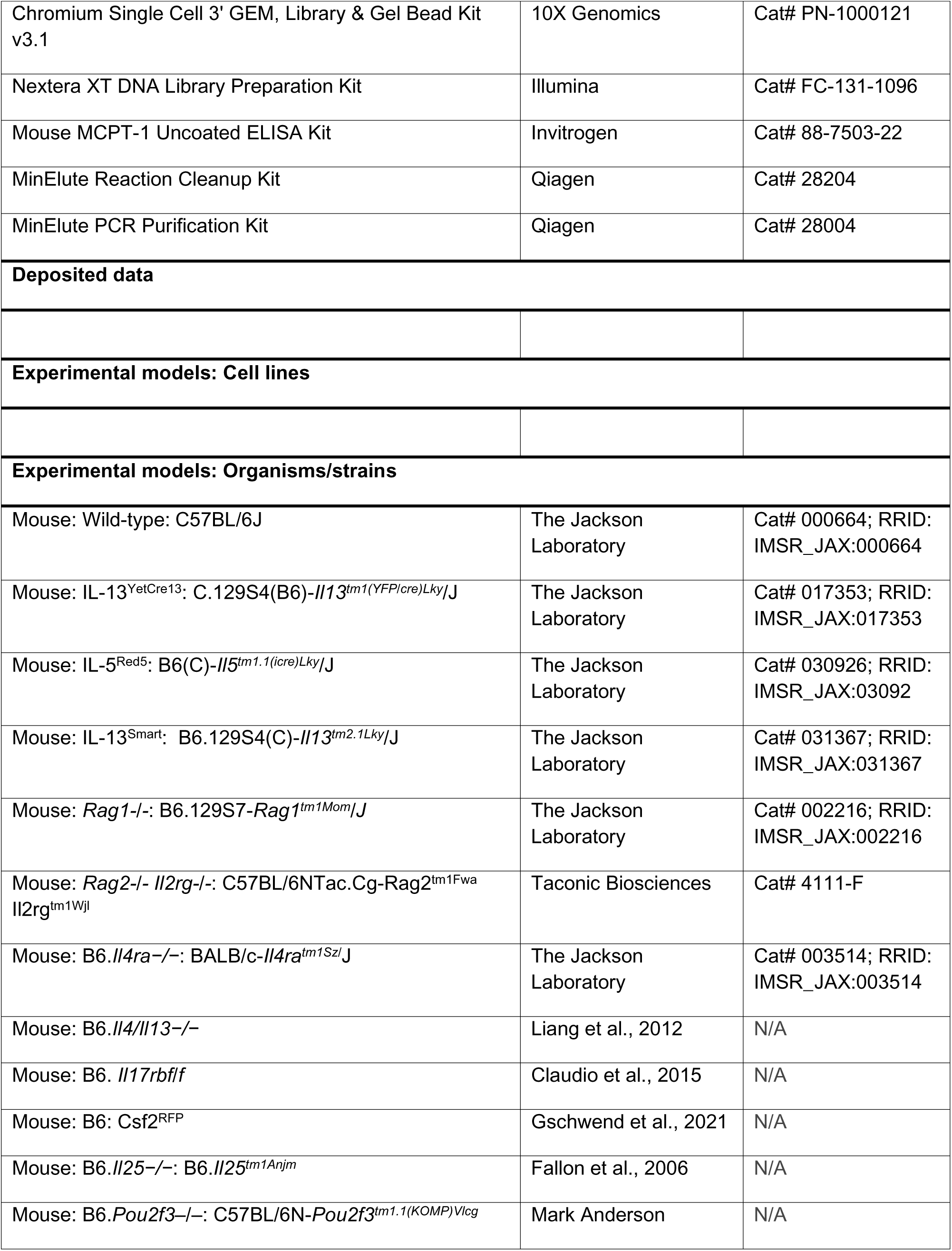
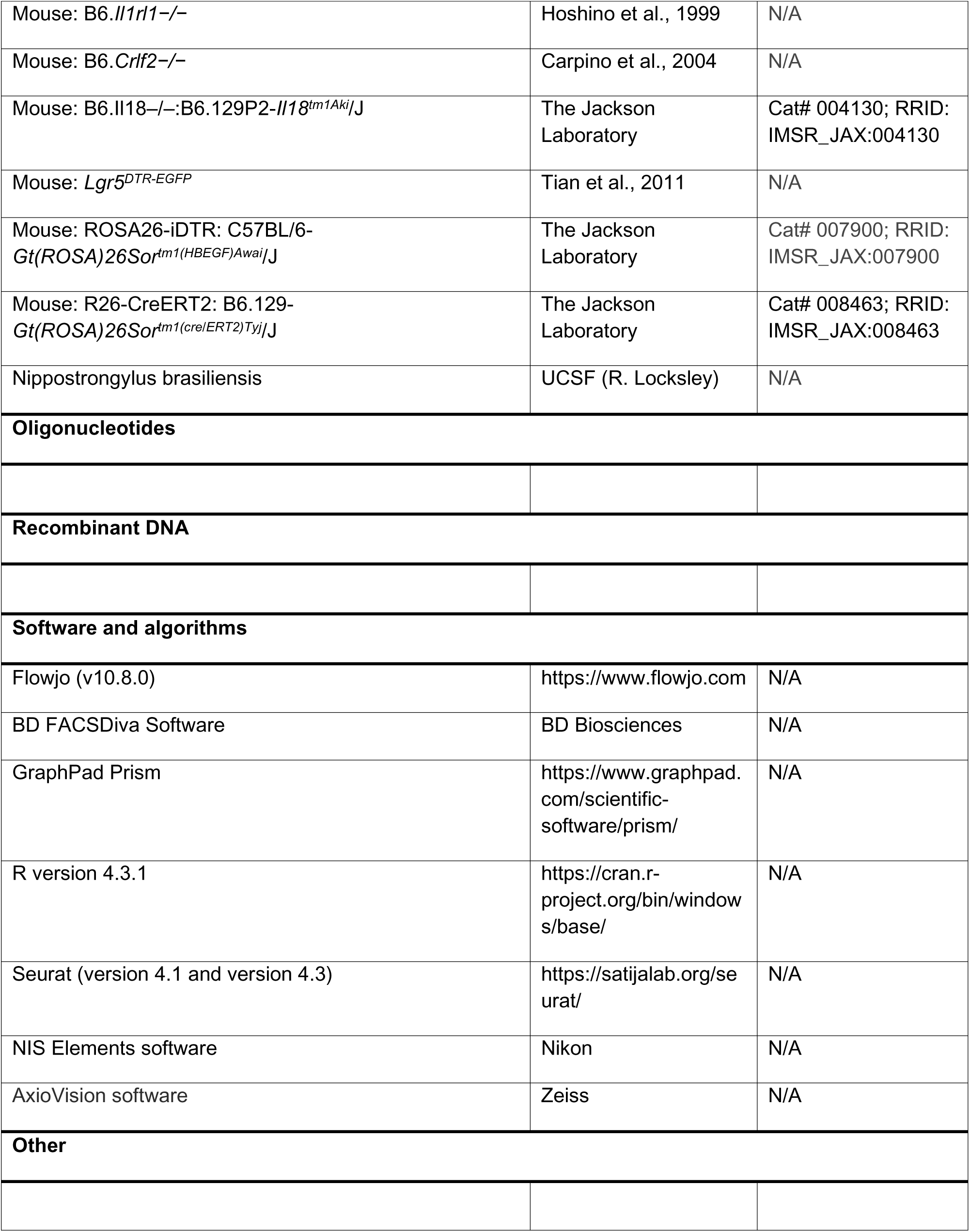

